# Genome replication dynamics of a bacteriophage and its satellite reveal strategies for parasitism and viral restriction

**DOI:** 10.1101/639039

**Authors:** Zachary K Barth, Tania V Silvas, Angus Angermeyer, Kimberley D Seed

**Affiliations:** Department of Plant and Microbial Biology, University of California, Berkeley, Berkeley, CA 94720, USA; Chan Zuckerberg Biohub, San Francisco, CA 94158, USA

## Abstract

Phage-inducible chromosomal island-like elements (PLEs) are bacteriophage satellites found in *Vibrio cholerae*. PLEs parasitize the lytic phage ICP1, excising from the bacterial chromosome, replicating, and mobilizing to new host cells following cell lysis. PLEs protect their host cell populations by completely restricting the production of ICP1 progeny. Previously, it was found that ICP1 replication was reduced during PLE(+) infection. Despite robustly replicating its genome, PLE produces relatively few transducing units, leading us to investigate if PLE DNA replication itself is antagonistic to ICP1 replication. Here we identify key constituents of PLE replication and assess their role in interference of ICP1. PLE encodes a RepA_N initiation factor that is sufficient to drive replication from the PLE origin of replication during ICP1 infection. In contrast to previously characterized bacteriophage satellites, expression of the PLE initiation factor was not sufficient for PLE replication in the absence of phage. Replication of PLE was necessary for interference of ICP1 DNA replication, but replication of a minimalized PLE replicon was not sufficient for ICP1 DNA replication interference. Despite restoration of ICP1 DNA replication, non-replicating PLE remained broadly inhibitory against ICP1. These results suggest that PLE DNA replication is one of multiple mechanisms contributing to ICP1 restriction.

## INTRODUCTION

Viral satellites are found in all domains of life and can have a profound impact on their helper viruses and their host cells (1–3). These sub-viral agents are known to worsen disease in humans (4) as well as plants (5), provide bacterial pathogens with toxins necessary for virulence (6), and serve as anti-viral immune systems in both single celled eukaryotes (7) and bacteria (8). As the parasites of viruses, satellites face distinct challenges in their lifecycles. Viruses typically need to subvert host cell nucleic acid metabolism in order to replicate their genome. In turn, viral satellites must find a way to subvert the subverters, so that the satellite’s genome can be replicated and mobilized in addition to, or to the exclusion of, the helper virus.

Within bacteria, four phylogenetically unrelated families of tailed-bacteriophage satellites have been discovered. These include satellite phage P4 and its relatives found in *Escherichia coli* (9, 10) the phage inducible chromosomal islands (PICIs) widespread throughout Firmicutes (11), the PICI-like elements (PLEs) found in epidemic isolates of *V. cholerae* (12), and the recently discovered Gram-negative PICIs found in Enterobacterialles and Pasturellales (13). Certain details of the life cycles of PLEs and their helper phage, ICP1, distinguish PLEs from other bacteriophage satellites. Both P4 and the well characterized subfamily of PICIs referred to as staphylococcal pathogenicity islands (SaPIs) confer partial restriction of their helper phages (14, 15). In contrast, PLEs completely restrict ICP1 production when they are able to progress through their replication cycle (12). This allows PLEs to function as effective abortive infection systems: individual ICP1 infected cells die, but since no phage are produced, the population as a whole is protected (12). PLEs’ more severe restriction of its helper phage likely relates to ICP1’s life cycle. ICP1 is only known to produce lytic infections that kill the host cell (16). In contrast, both P4 and PICIs parasitize temperate phages which occasionally integrate into the genomes of the cells they infect. For P4 and PICIs, it is not uncommon to find a helper phage and its satellite lysogenizing the same strain (9, 17). Since satellites rely on their helpers for mobilization, there can be intrinsic benefits to a low level of helper phage production that allows for co-lysogeny. If ICP1 kills every cell it can infect, cells that are potential hosts for PLEs, then it is to the PLEs’ benefit to completely restrict the production of infectious ICP1 progeny.

PLE’s use of ICP1 as a helper virus also has implications for PLE’s genome replication strategy. P4’s helper phage is known to rely on host encoded machinery (18), and while the replication of PICI helper phages have not been extensively characterized, comparative genomics suggest that some of the characterized PICI helpers hijack host cell replication machinery (19, 20). Similar to their helpers, both P4 and PICIs must redirect cellular machinery to the satellite genomes (21–23). Like well-studied lytic phages however, ICP1 encodes its own replication machinery (16). PLEs must therefore use a separate DNA polymerase from ICP1, or hijack ICP1’s DNA polymerase for their own replication. Either possibility provides a novel twist to bacteriophage satellite DNA replication. To excise PLE from the bacterial chromosome, the PLE integrase requires an ICP1-encoded recombination directionality factor (24), and PLE also requires the same viral receptor as ICP1 for transduction (12). Reliance on the same cell receptor as ICP1 suggests that PLE is packaged into ICP1 capsids just as P4 and PICIs are packaged into the capsids of their helper phage (2). PLE replication through use of ICP1’s replisome would then be in line with PLE’s reliance on ICP1 for multiple steps in the PLE lifecycle.

PLE’s severe parasitism of ICP1 has necessitated ICP1’s evolution of counter defenses. ICP1 host range on different PLEs varies among ICP1 isolates in a manner reminiscent of host-parasite co-evolution (12). So far, five distinct PLEs have been identified, and there has been a temporal succession of these elements, with a new PLE emerging around the same time as the previous PLE disappears from sequenced isolates. PLEs are prevalent, occurring in ~25% of *V. cholerae* isolates spanning a 60 year collection period (12). PLE(+) *V. cholerae* have been isolated from cholera patient stool samples alongside ICP1, suggesting that ICP1 infection, and PLE parasitism of ICP1, takes place within human hosts (8, 12, 24, 25). ICP1 isolates appear to have multiple strategies to overcome PLE (12), but the only mechanism identified so far is a phage-encoded CRISPR-Cas system (8, 25). Among PLE genes, only the PLE integrase has a recognized function (24), and the precise mechanism(s) by which PLEs restrict ICP1 continue to elude.

Given the crucial role of genome replication in viral propagation, the interface of PLE and ICP1 DNA replication is likely tied to PLEs’ ability to restrict ICP1. Previous work showed that PLEs can replicate upwards of 1000-fold following ICP1 infection (12) (Figure 1). This increase in PLE copy is accompanied by a 3 to 4-fold inhibition of ICP1 DNA replication. Curiously, PLEs do not transduce well under laboratory conditions, producing fewer than one PLE transducing unit per infected cell. Further, in these laboratory conditions, four of the five PLEs integrate seemingly randomly into one of *V. cholerae’s* many *Vibrio cholerae*-repeats (VCRs), but for PLE(+) *V. cholerae* isolates from nature, these four PLEs always occupy the same VCR, indicating that horizontal transmission may be rare (12). This suggests that transduction may play a minor role in the PLE lifecycle, and/or that it may be infrequent. The discrepancy between robust PLE replication and poor PLE mobilization led us to investigate the requirements for PLE replication, and whether PLE may bolster its anti-phage activity through increasing its copy number.

**Figure 1.**
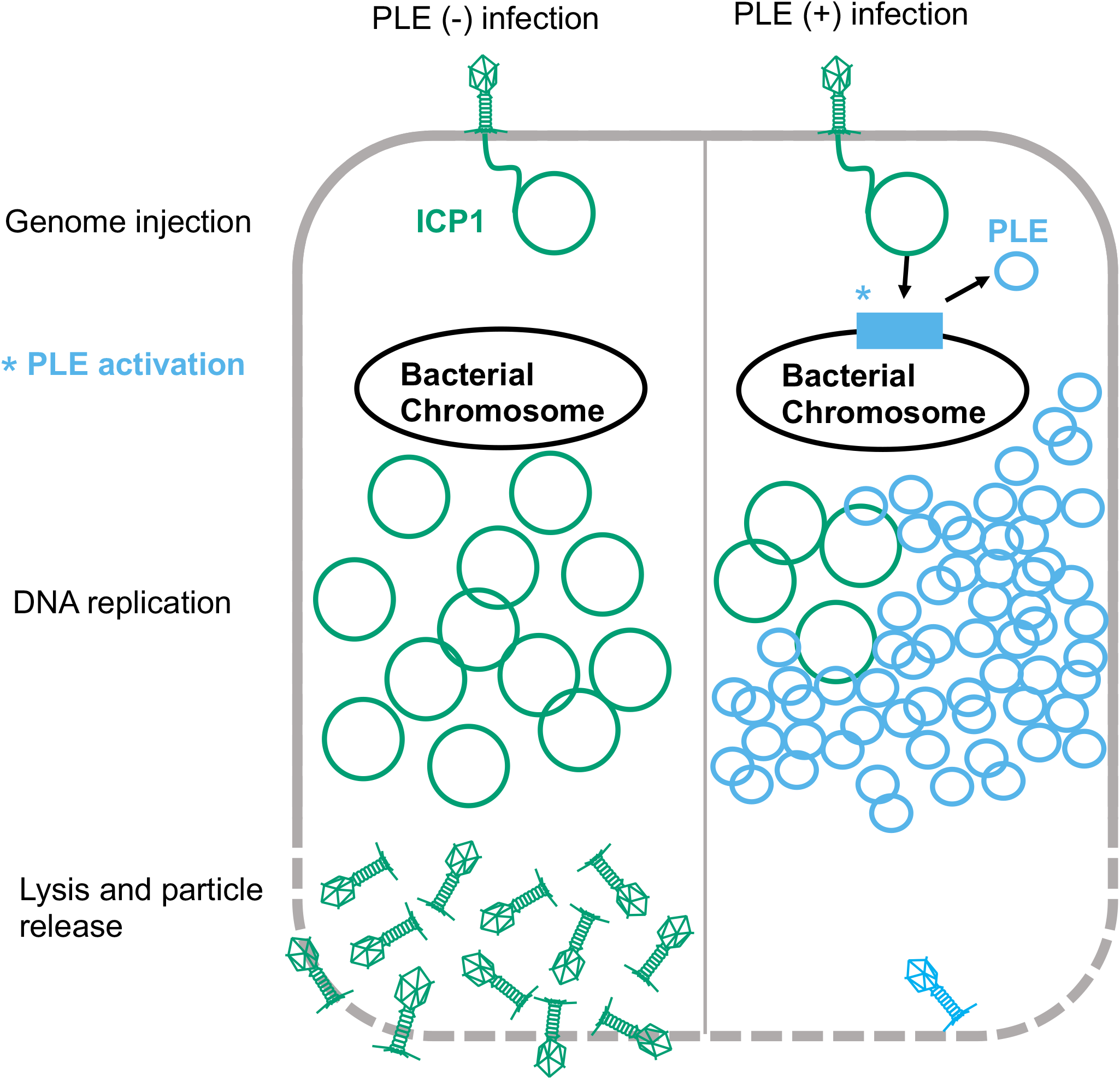
Model of ICP1 infection in PLE(-) and PLE(+) *V. cholerae*. ICP1 injects its DNA into *V. cholerae*; prior to DNA replication, ICP1 activity leads to PLE activation and excision. ICP1 DNA replication is reduced in the PLE(+) cell where the PLE replicates to high copy. Finally, the cell lyses and releases infectious particles. No ICP1 particles and a low number of PLE transducing particles are released from the PLE(+) cell.

Here, we define the replicon of PLE 1 (hereafter referred to as PLE), identifying an origin of replication and a PLE encoded replication initiation factor. The PLE replication initiator belongs to the RepA_N family of proteins, and to our knowledge is the first RepA_N protein functionally characterized in a Gram-negative bacterium. While PLE replication is neither necessary nor sufficient to provide anti-phage immunity against ICP1, loss of PLE replication does reverse the inhibition of ICP1 replication, and allows for a low level of ICP1 virion production, suggesting that PLE replication may be one of perhaps several parallel mechanisms that work in tandem to restrict ICP1.

## MATERIAL AND METHODS

### Strains and culture conditions

*V. cholerae* strains used in this study are derived from E7946. Bacteria were routinely grown on LB agar plates and in LB broth with aeration at 37°C. Antibiotics were supplemented as appropriate at the following concentrations: 75 μg/ml kanamycin, 100 μg/ml spectinomycin, 1.25 or 2.5 μg/ml chloramphenicol (*V. cholerae* for broth or plate conditions, respectively), 25 μg/ml chloramphenicol (*E. coli*), 100 μg/ml streptomycin. A detailed list of all strains used throughout this study can be found in Supplementary Table S1. ICP1_2006_E engineered to lack CRISPR-Cas (ΔCRISPR, Δcas2-3) (24) was used for all experiments. Phage susceptibility was determined using a soft agar overlay method wherein ICP1 was allowed to adsorb to *V.cholerae* for 10 minutes at room temperature before the mixture was added to molten LB soft Agar (0.3%) and poured onto 100mm x 15mm LB Agar plates. Plaques were counted after overnight incubation at 37°C. Efficiency of plaquing on mutant PLE strains was determined by dividing the phage titer obtained on the mutant PLE(+) strain by the phage titer obtained on PLE(-) strain.

### Generation of mutant strains and constructs

*V. cholerae* mutants were generated through natural transformation or *sacB* counter selection. Natural transformation was performed as described previously (26). For gene knockouts, splicing by overlap extension (SOE) PCR was used to generate deletion constructs with a spectinomycin resistance cassette flanked by frt recombination sites. Following selection of spectinomycin resistant mutants, a plasmid bearing an IPTG inducible Flp recombinase was mated into transformants and Flp expression was induced to generate in-frame deletions. The plasmid was cured by growing mutants under inducing conditions with 300μg/mL streptomycin. For plasmid expression constructs, a derivative of the pMMB67EH vector with a theophylline controlled riboswitch was used as previously described (24). For strains made via *sacB* counter selection, a marker-less deletion construct was generated using SOE PCR, and cloned into a pCVD442 suicide vector bearing the *sacB* counter selectable marker and an ampicillin resistance marker via Gibson assembly. LB Agar 10% sucrose plates were used to select for *sacB* loss and recombination of the mutant allele. All constructs were confirmed with DNA sequencing over the region of interest and primer sequences are available upon request.

### Real-time quantitative PCR

qPCR experiments were performed as previously described (12). Briefly, liquid cultures were infected with ICP1 at a Multiplicity of Infection (MOI) of 2.5 at OD_600_= 0.3. Samples were taken at 0 and 20 minutes post-infection, and boiled before serving as templates for IQ SYBR qPCR reactions. For assays involving induction of *repA*, 2mL cultures were grown with 1.25 μg/mL chloramphenicol for plasmid maintenance and induced for 20 minutes prior to infection using a final concentration of 1.5mM theophylline and 1mM IPTG starting at OD_600_= 0.17. Primers used for qPCR are listed in Supplementary Table S2.

### Protein purification

*E. coli* BL21 cells containing a pE-SUMO fusion to *repA* were grown to OD_600_= 0.5 at 37°C and induced with IPTG to a final concentration of 0.5mM. The cultures were then shifted to 16°C and grown for 24 hours. Cells were centrifuged and resuspended in lysis buffer (50mM Tris-HCl pH 8, 200 mM NaCl, 1mM BME, 0.5% Triton-X, 50mM imidazole, 1 Pierce™ Protease Inhibitor Mini Tablet (Thermo Scientific) and sonicated. Cell debris was removed by centrifugation (29,097 x g for 40 minutes), and the lysate was applied to a nickel resin affinity column (HisPur Ni-NTA Resin, Thermo Scientific). The column was washed with two column volumes of wash buffer (50mM Tris-HCl pH 8, 200mM NaCl, 1mM BME, 50mM imidazole), one column volume of an additional high salt wash (50mM Tris-HCl pH 8, 2M NaCl, 1mM BME, 50mM imidazole) to remove any residual DNA, and then eluted with elution buffer (50mM Tris-HCl pH 8, 200 mM NaCl, 1mM BME, 300mM imidazole). The eluate was dialyzed with sizing buffer (50mM Tris-HCl pH 7.5, 150mM NaCl, 1mM Dithiothreitol) in 10K MWCO SnakeSkin dialysis tubing (Thermo Scientific), and the His6-Sumo-tag was cleaved with 1 μL SUMO protease per 100 μg of protein. The eluate was fractionated on a HiLoad 16/60 Superdex 75 size-exclusion column (GE Healthcare) and fractions were analyzed using SDS page. Protein was concentrated using a Amicon Ultra 15mL 3K NMWL centrifugal filter (Millipore Sigma).

### Electrophoretic mobility shift assays (EMSAs)

dsDNA probes used for RepA DNA binding experiments were generated using PCR with primers listed in Supplementary Table S2. 50 fmol of probe was incubated with purified RepA at 30°C for 20 minutes in 20μL reactions with 10mM Tris pH 7.8, 10% Glycerol, 1μM TCEP, 10mM MgCl_2_, and 0.4 μg poly d(I-C) (Sigma-Aldrich) serving as a nonspecific competitor. The full reaction volume was then loaded onto 2.5% agarose 0.5X Tris-borate gels with 1x Thiazole Green, and ran for 20 minutes at 120V before visualization.

### Preparation of phage infection samples for DNA sequencing

A 6mL bacterial culture was infected at OD_600_= 0.3 with ICP1 at an MOI of 1. At the indicated time points, 1mL was removed from the culture tube and mixed with 1mL ice cold methanol to stop DNA synthesis. These samples were pelleted at 21,694 × g for 2 minutes at 4°C. The methanol was removed through aspiration and the pellet was washed with 1 mL cold Phosphate-buffered saline. Pellets were frozen in liquid nitrogen and stored at −80°C until total DNA was isolated using the QIAGEN DNeasy Blood and Tissue Kit. Sequencing libraries were prepared using NEBNext® Ultra™ II FS DNA Library Prep Kit. Paired-end sequencing (2 × 150 bp) was performed on an Illumina HiSeq4000 (University of California, Berkeley QB3 Core Facility).

### DNA-seq reads mapping

Illumina sequencing reads for each timepoint were mapped to the appropriate reference sequence using Bowtie 2 v2.3.4.1 (27) with default settings except for the following: ‘--end-to-end’ and ‘--very-sensitive’. Mapping files were sorted and indexed with samtools v1.5 and binned with breseq BAM2COV v0.33.0 (28): ICP1 1000bp, PLE 150bp. Plots were generated in Python3 with the matplotlib module v3.0.3 (29). For plotting the abundance of a specific genome in a sample, the genome per million (GPM) was calculated. GPM is calculated in the same manner as the previously described transcript per million (TPM) (30).

### Protein structure visualization

Protein structure figures were generated using PyMOL (The PyMOL Molecular Graphics System, Version 2.0 Schrödinger, LLC). Protein structure alignments were generated using the cealign command (31). The electrostatic distribution was determined and visualized using the PDB2PQR server, and the APBS plugin for PyMOL (32).

### Transduction assays

Transduction assays were performed as previously described (12). Briefly, donors were grown to OD_600_= 0.3 and infected with ICP1 at an MOI of 5. Cultures were incubated for 5 minutes before being washed with fresh LB to remove unbound phage. The infected cultures were incubated for 30 minutes, and 100μL lysate was added to 100μL of an overnight culture of the recipient strain. The mixture was incubated for 1 hour at 37°C with aeration before plating on selective media.

## RESULTS

### PLE alters and diminishes ICP1 replication

PLE was previously shown to replicate to high copy during ICP1 infection and reduce ICP1 DNA replication compared to a PLE(-) control (12) (Figure 1), however, these results were obtained through qPCR and only assessed a single ~100bp target sequence. To obtain a more complete understanding of PLE replication dynamics and the PLE’s impact on ICP1 replication kinetics, we performed deep sequencing of total DNA during an ICP1 infection time course using PLE(-) and PLE(+) *V. cholerae*. ICP1 produces new progeny virions by 20 minutes post-infection in PLE(-) cultures, and PLE(+) cultures lyse 20 minutes post-infection (12), therefore to evaluate total DNA content in infected cells at early, middle and late time points (while avoiding potential DNA loss due to lysis), we collected samples at 4, 8, 12 and 16 minutes post-infection. Samples were prepped for total DNA at each time point and the resulting Illumina sequencing reads were mapped against the *V. cholerae*, ICP1, and PLE genomes. Consistent with the anticipated rapid kinetics of ICP1 infection in PLE(-) *V. cholerae*, the abundance of ICP1 reads increased within 8 minutes post-infection and ICP1 DNA comprised roughly half of the total DNA content by 16 minutes post-infection (Supplementary Table S3). To account for the relatively small size of the ICP1 genome compared to the *V. cholerae* chromosomes, we normalized the reads mapped per element to element length and the total reads per sample to determine the genomes per million (GPM) of each entity in the samples. We observed that ICP1 DNA replication robustly overtakes the cell and phage genomes are more abundant than copies of the *V. cholerae* large and small chromosomes by 12 minutes post-infection (Figure 2A). In contrast, ICP1 DNA replication is less robust in the presence of PLE. Specifically, the proportional abundance of ICP1 DNA is relatively unchanged at 4 and 8 minutes post-infection of PLE(+) cells, but ICP1 DNA replication begins to dramatically lag by 12 minutes post-infection compared to PLE(-) infection (Supplementary Table S3). In the PLE(-) condition, ICP1 relative reads abundance doubles from roughly one quarter to one half of total reads between 12 and 16 minutes post-infection, while in the PLE(+) condition ICP1 abundance increases very little between 12 and 16 minutes post-infection (Supplementary Table S3). The defect observed in ICP1 replication correlates with PLE’s own robust replication. By 8 minutes post-infection, PLE is already the most abundant element in terms of copy number (Figure 2B). Between 8 and 16 minutes post-infection, the abundance of PLE DNA grows to comprise approximately 19% of total reads, overtaking ICP1 in total DNA (Supplementary Table S3). In terms of genome copy at 16 minutes post-infection, PLE outnumbers ICP1 approximately eight-fold (Figure 2B). The temporal dynamics of PLE and ICP1 DNA replication support the notion that interference of ICP1 replication may be linked to the PLE’s own replication.

**Figure 2.**
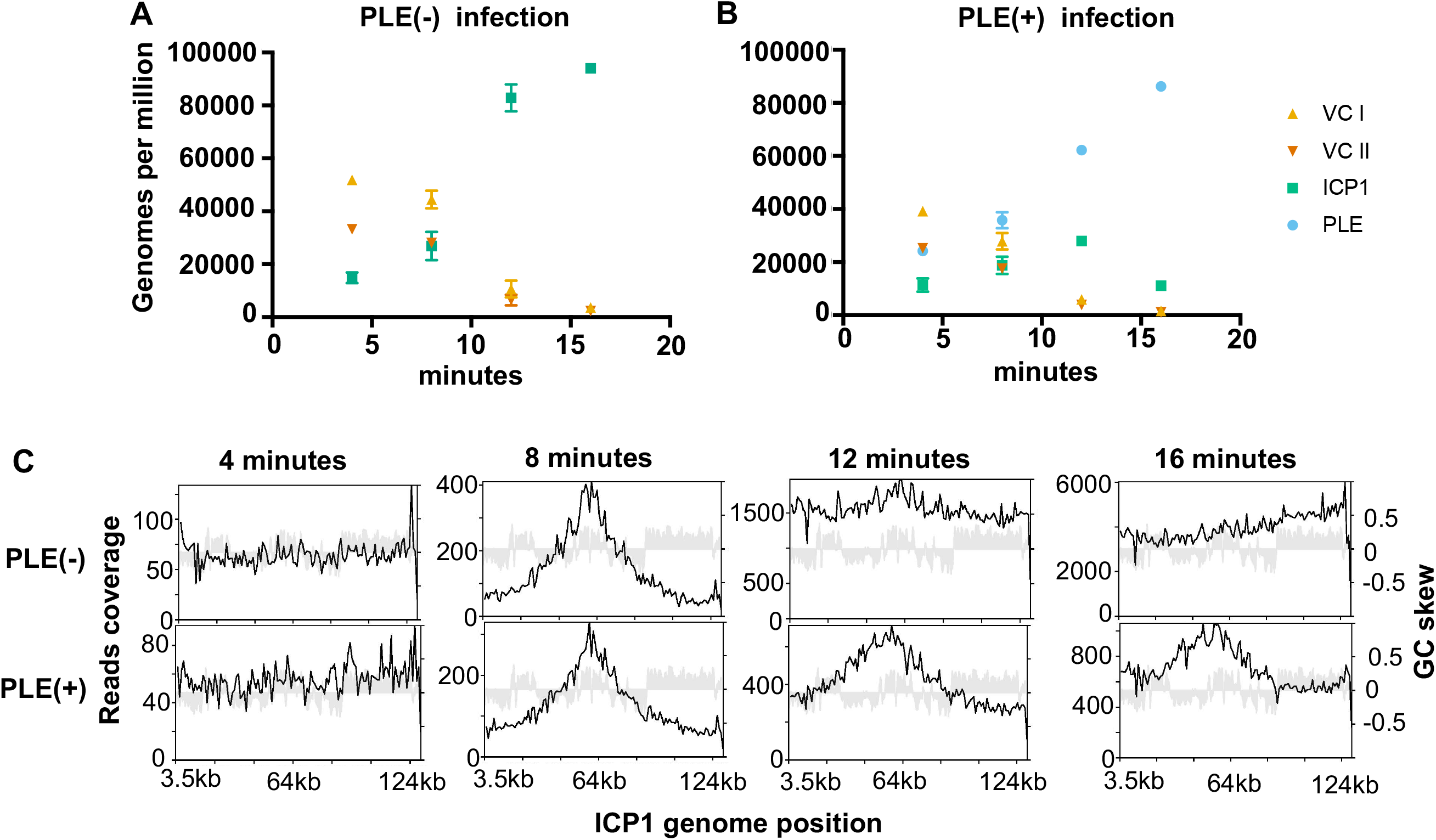
PLE robustly replicates following infection while altering ICP1 replication. (**A** and **B**) Genomes per million (GPM) of total DNA mapping to the *V. cholerae* large (VC I) and small (VC II) chromosomes, ICP1, and the PLE across an infection time course in PLE(-) (**A**) and PLE(+) (**B**) *V. cholerae*. Samples were taken at 4, 8, 12, and 16 minutes post infection, data show the average and standard deviation of three independent experiments. (**C**) Reads coverage plots across the ICP1 genome for PLE(-) (top) and PLE(+) (bottom) during infection. Data from one replicate are shown, replicate experiments are shown in Supplementary Figures S1 and S2.

In addition to monitoring the relative changes in abundance of discrete genetic elements during phage infection, we evaluated the profiles of sequence coverage across ICP1 and PLE genomes (Figure 2C and Supplementary Figures S1-S3) While the distribution of reads across the ICP1 genome was similar in PLE(+) and PLE(-) conditions at 8 minutes post-infection, ICP1’s coverage profile was markedly different at 12 and 16 minutes post-infection between the two conditions (Figure 2C). At 8 minutes post-infection, a peak in ICP1 reads can be seen near the 60kb position, and reads abundance decreases with increasing distance from that point. The observed pattern in ICP1, which is present in both PLE(+) and PLE(-) infections, is consistent with the predicted coverage of an element that replicates bidirectionally through theta-replication from a single origin of replication (ori) (33). At 16 minutes post-infection in the PLE(-) condition the peak reads abundance was shifted to one end of ICP1’s genome (Figure 2C). These results suggest activation of an additional ICP1 ori late in infection. Additionally, the distribution of reads decreases gradually in the upstream direction from the peak, and sharply drops downstream of the peak. Such a distribution is suggestive of a rolling circle mode of replication (33), which is consistent with a number of phages that are known to transition to rolling circle late in infection (34). By contrast, at 16 minutes post-infection in PLE(+) *V. cholerae*, the profile of ICP1 reads more strongly resembled the profile at 8 minutes post-infection than it did to the coverage profile 16 minutes post-infection in the PLE(-) condition. The change in ICP1 reads distribution suggests that PLE might alter ICP1 replication origin choice and impair the progression from theta to rolling circle replication.

Intriguingly, the reads peak at near the end of ICP1’s annotated genome was prominently visible at 4 minutes post infection, before ICP1 replication has taken place (Figure 2C). We speculated that this reads peak corresponds with the terminus of infecting ICP1 particles prior to genome circularization as it has previously been established that termini can lead to sequencing biases following DNA library preparation (35). We found that this reads bias was also present in DNA from purified phage particles (Supplementary Figure S4). PhageTerm, which is designed to identify phage termini and packaging methods (35), identified this reads peak as a packaging (pac) site and predicted a headful packaging mechanism for ICP1. The terminus is located in a 1.3kb orfless space between *gp1* and *gp2* (Supplementary Figure S4A and B). PhageTerm predicts the location of the pac site at 431bp on the annotated (+) strand, and 891bp on the annotated (-) strand (Supplementary Figure S4C). The loss of this peak by 8 minutes post-infection (Figure 2C) likely reflects circularization of the phage genome following cell entry. Together, ICP1’s changing coverage profile over the course of infection, and PhageTerm analysis of phage particle DNA suggests that rolling circle initiation and genome packaging may be linked for ICP1. That the shift in ICP1 coverage profile was profoundly reduced in the PLE(+) background suggests that PLE interferes with the rolling circle mode of ICP1 replication, and that this interference may perturb later steps (i.e. DNA packaging) necessary for ICP1 to complete its lifecycle.

### PLE encodes its own replication initiator, but does not replicate autonomously

To better understand the relationship between PLE and ICP1 DNA replication, we next sought to identify the constituents of the PLE replicon. The PLE genome is 18kb and organized into multiple predicted gene clusters (Figure 3A). Between PLE *orf5* and *orf7*, there is a 2.7kb non-coding region (NCR) which has four repeat sequences (Figure 3B, Supplementary Figure S5).

**Figure 3.**
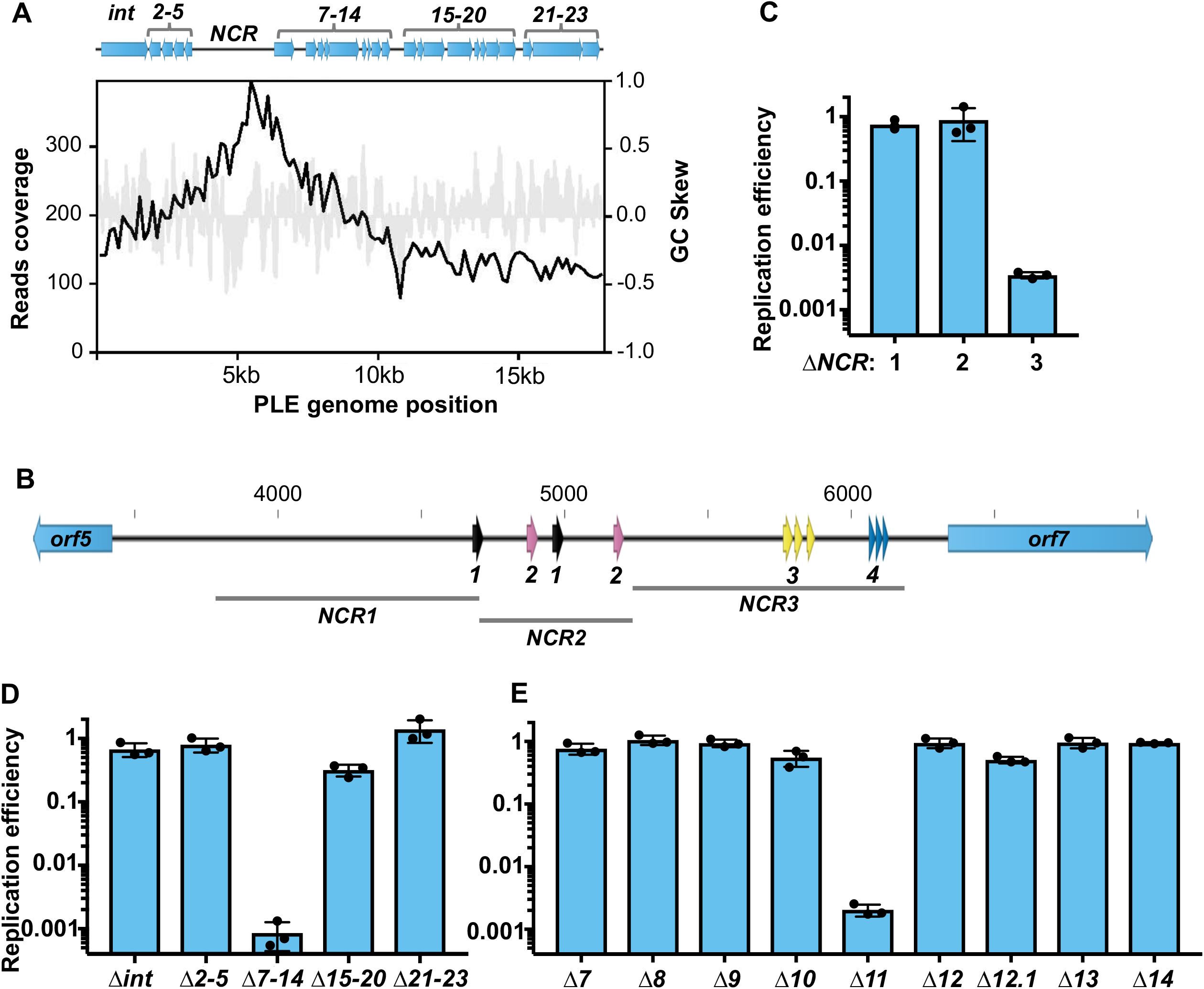
A single PLE-encoded orf and a noncoding region are necessary for PLE replication. **(A)** A representation of the PLE genome (top) with coverage 8 minutes post ICP1 infection plotted below for a representative sample. Gene clusters mutated for analysis are labelled. PLE coverage for additional replicates at all time points is shown in Supplementary Figure S3. **(B)** PLE’s noncoding region (NCR) between *orf5* and *orf7*, with repeat sequences shown as arrows. Repeat sequences share colors for each repeat type, and are designated as repeats 1, 2, 3, or 4. Regions of the NCR deleted for analysis in (C) are shown. Panels **C-E**: replication of PLE mutants 20 minutes post-infection with ICP1 as assessed by qPCR. Replication efficiency is relative to a WT PLE control. **(C)** Replication of Δ*NCR* mutants. **(D)** Replication of PLE gene cluster knockouts. **(E)** Replication of individual gene knockouts of the orfs contained in cluster *7-14*.

Frequently, repetitive sequences serve as binding sites for replication machinery at phage and plasmid origins of replication (36). Within bacterial genomes, there is also a bias for coding sequence in the leading strand (37), and this is consistent with an ori being between divergently transcribed operons. These features led us to hypothesize that the PLE NCR serves a function in replication. This was further evidenced by PLE’s replication reads profile, which showed a peak approximately 1kb upstream of *orf7* at 8 minutes post-infection, suggesting that the PLE ori is located in the NCR (Figure 3A). To test if the NCR contained sequence necessary for PLE replication, we generated three strains designated *NCR1*, *NCR2* and *NCR3* that each possessed a 0.5-1kb deletion within the NCR excluding predicted promoters for *orf5* and *orf7* (Figure 3B). Following ICP1 infection we found that *NCR1* and *NCR2* were dispensable for PLE replication, however, *NCR3*, containing repeat sequences 3 and 4, was necessary for replication (Figure 3C). This suggested that the PLE ori was contained within *NCR3*, and that repeat 3 and repeat 4 may be involved in replisome recruitment. We next sought to determine whether any predicted PLE open reading frames (ORFs) are necessary for PLE replication. We began by screening PLE gene cluster knockouts (24) and we observed that one gene cluster, containing *orf7* through *orf14*, was necessary for PLE replication (Figure 3D). To identify the ORF responsible, we constructed individual gene knockouts within this cluster and screened for replication defects during ICP1 infection. We found that a single open reading frame, *orf11* (https://www.ncbi.nlm.nih.gov/protein/AGG09405.1/), was necessary for PLE replication (Figure 3E). Given the requirement of *orf11* for PLE replication, and further analyses supporting its designation as a replication initiation protein (discussed below) we designated *orf11* as *repA*.

In both P4 and SaPIs, satellite replication is autonomous following the satellite’s transcriptional activation by the helper phage (21, 38). Having determined that PLE-encoded *repA* is necessary for PLE replication, we sought to elucidate if expression of *orf11* was sufficient to drive autonomous replication of PLE. We complemented PLE Δ*rep*A with ectopically expressed *rep*A and measured PLE copy number increase in the presence and absence of phage. RepA expression was able to drive PLE replication, but only in cells infected by ICP1 (Figure 4A). To rule out the possibility that PLE replication requires additional redundant PLE genes activated by ICP1 that may have been missed in our genetic screen, we next set out to define the minimal unit required for PLE replication. Previous work showed that ICP1 infection triggers excision of a ‘miniPLE’, consisting of the PLE encoded integrase together with a kanamycin resistance marker flanked by the PLE attachment (att)-sites (24). We built on this existing platform and constructed a ‘midiPLE’ which differs only by the presence of the NCR containing the PLE ori on the self-excising miniPLE (Figure 4B). The midiPLE replicated to a substantial level that depended upon ICP1 infection and *repA* expression, though it was able to reach only approximately one fourth that of complemented PLE Δ*repA*. This result confirmed that *repA* and the PLE noncoding region are sufficient to drive PLE replication, but only in the presence of ICP1. PLE’s lack of autonomous replication is a striking contrast to previously characterized bacteriophage satellites. PLE’s reliance on ICP1 for replication and not just activation of gene expression suggests that PLE may parasitize ICP1-encoded gene products for replication, though it is also possible that PLE uses components of *V. cholerae’s* replication machinery that are expressed or activated only upon ICP1 infection.

**Figure 4.**
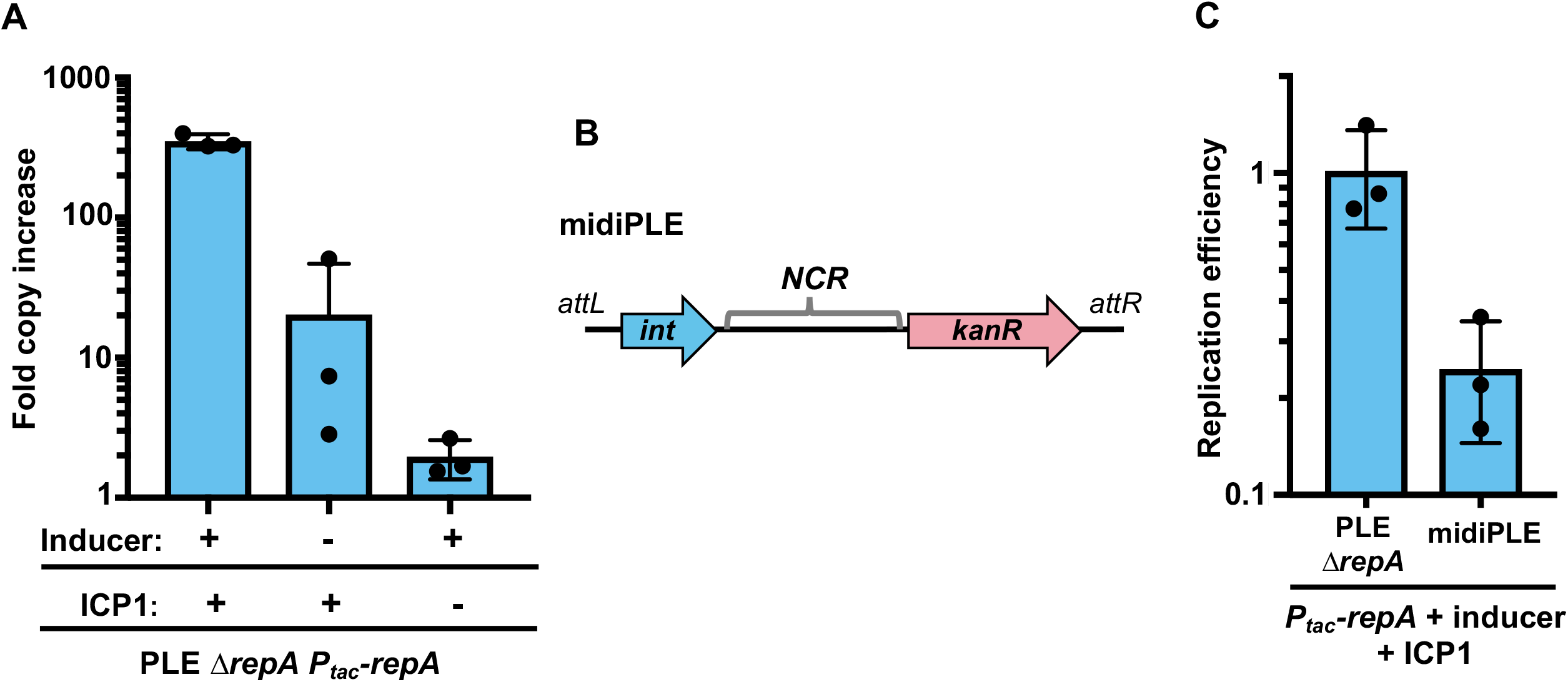
RepA drives PLE replication in the presence of ICP1. (**A**) Complementation of PLE Δ*repA*. PLE fold copy increase 20 minutes post-infection is shown in the presence of ICP1 and inducer, ICP1 alone, and inducer alone. **(B**) A diagram of the midiPLE construct used to assess the minimal requirements for PLE replication (not to scale). Attachment sites, the PLE integrase, and the noncoding region (NCR) are present along with a kanamycin resistance gene (*kanR*). (**C**) Replication of a RepA complemented Δ*repA* strain and midiPLE relative to a WT PLE control 20 minutes post ICP1 infection.

### The PLE encoded replication protein RepA resembles Gram-positive plasmid initiation factors

Although the structure and function of PLE’s RepA has not been previously elucidated, the x-ray crystal structure of the N-terminal domain (NTD) of RepA (RepA-NTD) has been solved and deposited in the Protein Data Bank (PDB ID: 4RO3) (Figure 5A). While primary sequence similarity (using BLASTP) is not evident, using HHpred (39), we found that PLE RepA-NTD has substantial structural similarity to the pKS41 and pTZ6162 plasmid RepA proteins from *Staphylococcus aureus*, as well as more distant similarity to the replication protein DnaD from *Bacilus subtilis* (PDB ID: 4PTA, 4PT7, and 2v79). Both of the *S. aureus* RepA proteins serve as replication initiators for plasmids coding for multidrug resistance and belong to the RepA_N family of plasmid replication proteins. The RepA_N protein family is comprised mostly of initiation factors for theta-replication of plasmids found mainly in the Firmicutes (40, 41). This protein family is named for the conservation of the NTD which structurally resembles the NTD of the Gram-positive primosome component DnaD (42). In the RepA_N family, the NTD mediates DNA binding, while the C-terminal domain (CTD) of these proteins appear to be specific to host genus, and may perform host specific functions (40). Indeed, PLE RepA’s CTD did not have any strong similarity to other characterized proteins detectable by HHpred or HHblits (expect >1 for all hits).

**Figure 5.**
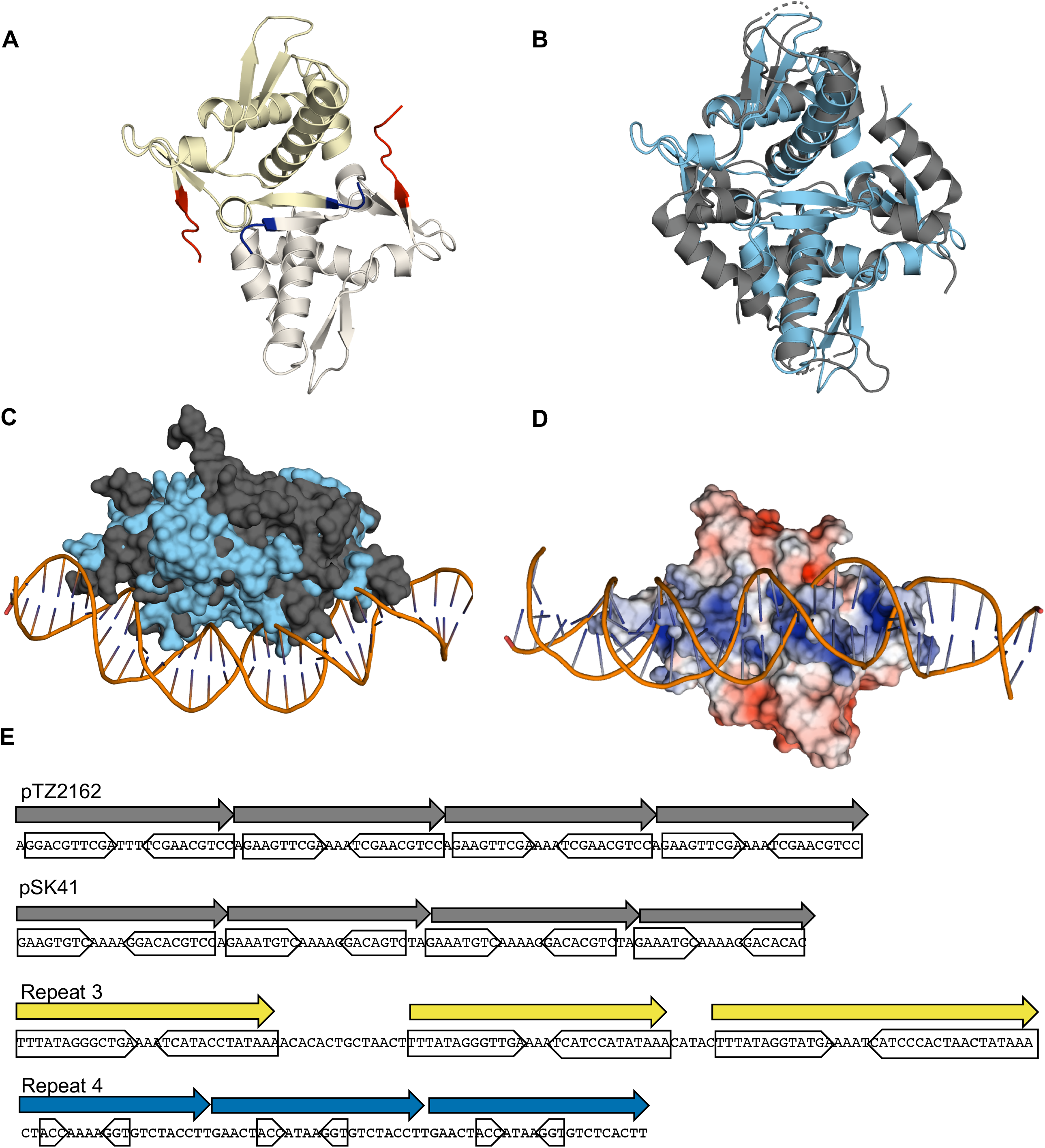
PLE RepA is a RepA_N family protein. (**A**) Cartoon representation of the PLE RepA-NTD dimer. Monomers are differently colored in yellow and white. The N and C termini of the monomers are colored blue and red respectively. (**B**) Alignment of the NTD dimers of PLE RepA and pTZ2162 RepA in light blue and dark grey, respectively, depicted in cartoon representations (RMSD = 4.527 over 176 residues). (**C**) Surface view of PLE RepA-NTD dimer in light blue aligned with pTZ2162 RepA-NTD dimer in dark gray bound to substrate DNA. (**D**) Electrostatic potential map, turned 90 degrees as (C), of PLE RepA-NTD dimer aligned to the pTZ2162 RepA-dsDNA bound structure. Positive (blue) and negative (red) charges are indicated on the surface. (**E**) Binding iterons for the RepA initiators of pTZ2162 and pSK41 are shown alongside repetitive sequences found in the putative PLE origin of replication. Direct repeats are denoted with an arrow, while the sequence comprising inverted sub-repeats is underlined. Sequence for the PLE (-) strand is shown to make the central poly-A tract apparent.

The PLE RepA-NTD structure aligns well with the crystal structures of the NTDs from *S. aureus* pTZ6162 and pSK41 RepA initiation factors, highlighting shared tertiary structure (Figure 5B, Supplementary Figure S6). Notably, all three proteins crystalized as dimers with monomers in the same orientation, suggesting a conserved dimer interface, and the potential for a conserved method for DNA binding. A crystal structure for the pTZ6162 RepA-NTD dimer bound to its cognate iteron dsDNA sequence has also been solved (PDB ID: 5kbj) (42),and we were able to align the PLE RepA-NTD to this structure (Figure 5C). In the original pTZ6162 structure, the surface of the protein that is bound to the DNA is electropositive (42), consistent with binding activity for the electronegative dsDNA sugar-phosphate backbone. The corresponding surface in the PLE RepA-NTD structure is electropositive as well, suggesting conserved DNA binding activity (Figure 5D). An electropositive DNA binding interface is also observed in the pSK41 RepA structure suggesting maintenance of a shared DNA binding region among these proteins (Supplementary Figure S7A-C). Notably, the DnaD-NTD is less electropositive along the corresponding surface, which is expected since the DnaD-NTD, despite its structural similarity to RepA_N-NTDs, does not bind DNA (40). Given PLE RepA’s structural similarity to the RepA_N family, its similar electrostatic profile, and its shared role as a replication factor for a mobile genetic element, we conclude that PLE RepA belongs to the RepA_N protein family.

Like other replication initiation factors (36), RepA_N family proteins bind to repetitive iteron sequences at their cognate ori. Most characterized RepA_iterons are semi-palindromic direct repeats, containing inverted repeats that converge on a poly-A tract (40) (Figure 5E). These same sequence features are apparent in repeat 3 and repeat 4 in *NCR3* (Figure 5E), which is necessary for PLE replication (Figure 3B). The iterons for pTZ2162 and pSK41 have repeats of 9 and 8bp respectively. Repeat 3 in PLE has inverted repeats that are longer at 13bp, while those in repeat 4 are only 3bp long. Most characterized RepA_N iteron inverted repeats are at least 5bp long, but obvious inverted repeats are not always discernable (40). To determine if PLE’s RepA is capable of binding to either of the repetitive sequences in *NRC3*, we purified RepA and assessed binding to the putative ori^PLE^ through an EMSA. We tested RepA binding using 141bp probes containing the repeat 3 sequence, the repeat 4 sequence, and part of *NCR2* containing repeats 1 and 2. Consistent with *NCR2* being unnecessary for replication, the probe corresponding to repeats 1 and 2 did not bind to RepA (Figure 6). For the probes from *NCR3*, repeat 3 but not repeat 4 was shifted on the gel, confirming that RepA binds the repeat 3 sequence. This is consistent with our genetic analysis that *NCR3* is required for PLE replication, and shows that repeat 3 serves as the binding site for the PLE replication initiator protein RepA.

**Figure 6.**
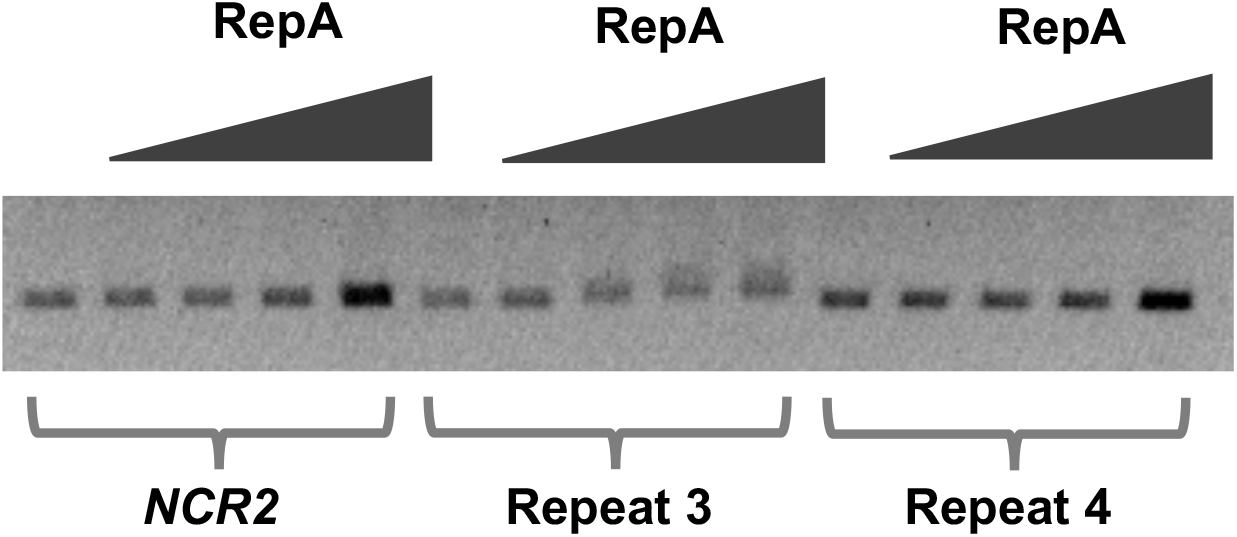
RepA binds to repeat 3 in the PLE noncoding region. An electrophoretic mobility shift assay using probes from the PLE noncoding region. RepA binding was tested for probes containing the repeat 3 or repeat 4 sequence from the PLE *NCR3*, as well as a probe matching the repeat 1 and repeat 2 sequences in *NCR2*. Additional replicates of this experiment are shown in Supplementary Figure S8.

### Non-replicating PLE alters ICP1 replication dynamics without lowering ICP1 genome copy

Having identified the necessary components of the PLE replicon, we sought to assess the importance of PLE replication for the PLE’s lifecycle and anti-phage activity. Following excision and replication, PLE can be transduced to recipient *V. cholerae* cells (12). We hypothesized that PLE replication would be necessary for its transduction, therefore we performed transduction assays, comparing Δ*repA* PLE complemented with either *repA* or an empty vector control. As expected, PLE transduction was below the limit of detection in Δ*repA* PLE complemented with an empty vector control (Supplementary Figure S9), and the transduction defect for Δ*repA* PLE could be complemented by *repA* in trans, restoring PLE transduction to levels near to wild-type (WT) PLE (3.8 × 10^4^ units/mL, (12)).

The finding that high PLE copy is needed to facilitate PLE transduction is intuitive, but under these laboratory conditions PLE produces fewer than one transducing unit per infected cell, despite PLE’s robust replication (12). This lead us to question if PLE replication contributes to PLE’s anti-phage activity. ICP1 replication is reduced in a PLE(+) infection (Figure 2). A potential mechanism of PLE impairment of ICP1 replication could be through the consumption of replication resources. Robust PLE replication might exhaust dNTP pools, and since PLE only replicates during ICP1 infection, PLE might also competitively restrict ICP1’s access to its own replisome. Therefore, we next tested ICP1 replication in non-replicating PLE strains using qPCR, and observed that ICP1 replication was restored to the levels seen in PLE(-) infection conditions (Figure 7A). This restoration led us to question if midiPLE replication could impair ICP1 replication simply by using up replication resources. During ectopic expression of *repA*, midiPLE did not impair ICP1 replication, while Δ*repA* PLE did (Figure 7B). Consistent with this result, we did not observe any defect in ICP1 plaque formation on *V. cholerae* harboring a replicating midiPLE (Supplementary Figure S10). This suggests that PLE replication reduces ICP1 copy through an independent mechanism, which may be dependent on PLE gene dosage increase, or by reaching a level of replication not achievable with the midiPLE.

**Figure 7.**
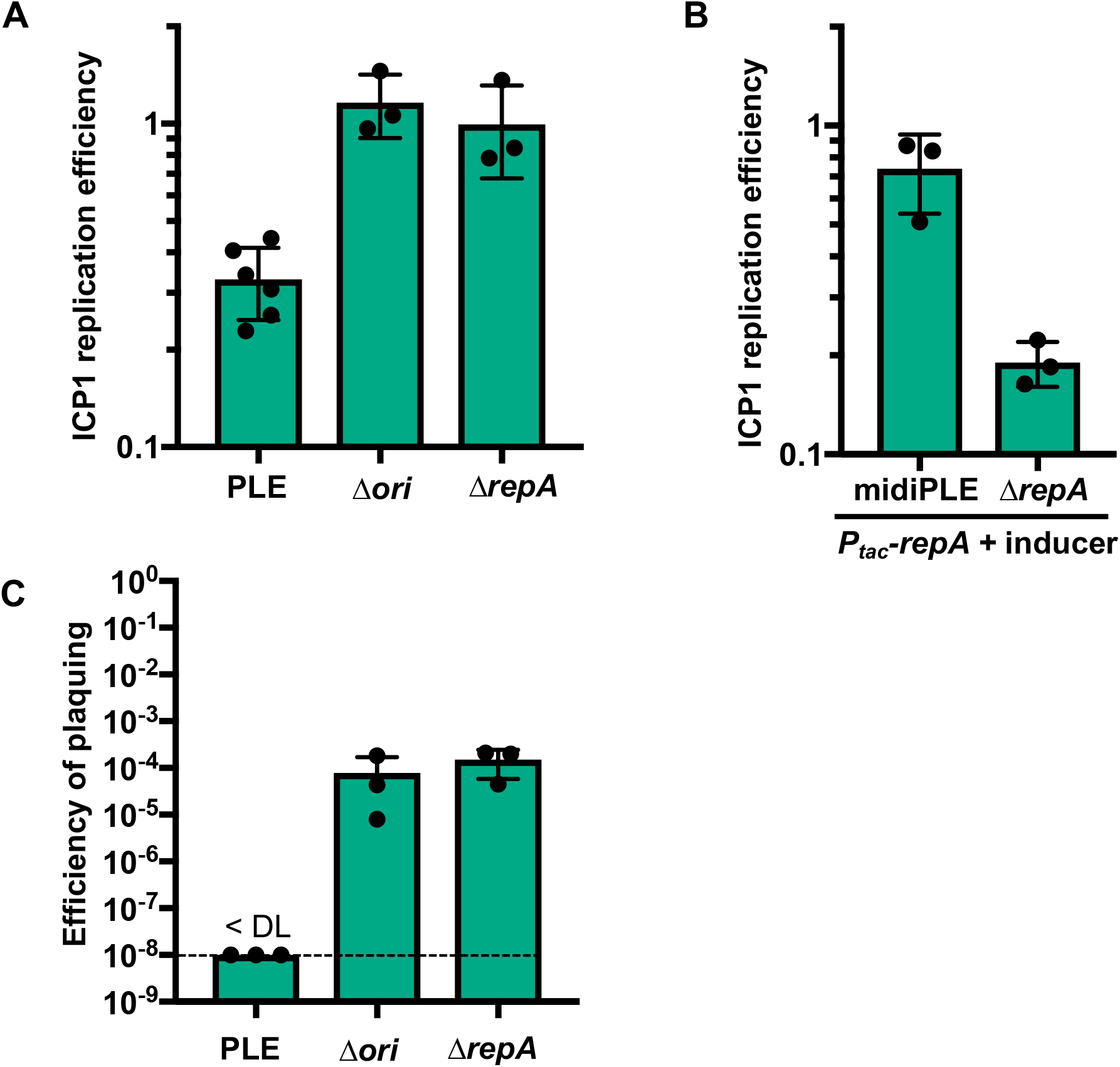
Loss of replication impairs PLE anti-phage activity. (**A**) ICP1 replication in WT and mutant PLE(+) strains as assessed by qPCR. Replication efficiency is relative to ICP1 infection of PLE(-) *V. cholerae* 20 minutes post-infection. (**B**) Replication of ICP1 as assessed by qPCR in RepA complemented midiPLE and Δ*repA* PLE infection relative to an un-complemented midiPLE control. (**C**) Efficiency of plaquing (EOP) for ICP1 on WT PLE and non-replicating PLE mutant hosts. EOP is relative to a PLE(-) control strain, and the limit of detection (DL) is 10^−8^.

The roughly four-fold decrease in ICP1 replication that occurs in PLE(+) cultures would not likely be sufficient for the complete restriction of ICP1 that is observed (12), but is likely to be a contributing mechanism. To investigate this, we performed ICP1 plaque assays on non-replicating PLE mutant hosts. The PLE Δ*repA* and Δ*ori* mutants still blocked plaque formation (data not shown), however, the mutants were more susceptible to ICP1 than WT PLE as some small plaques were visible high phage concentrations. Therefore we quantified ICP1’s efficiency of plaquing (EOP) on these non-replicating PLE mutants and observed a near 10,000-fold increase in EOP compared to WT PLE, on which phage do not form any detectable plaques (Figure 7C). This level of plaque formation is still 10,000-fold lower than the yield of a PLE(-) host, demonstrating that although PLE replication is not necessary for PLE mediated anti-phage activity, PLE replication bolsters or acts synergistically with other PLE encoded anti-phage activities.

We observed that PLE replication decreases the level of ICP1 replication and coverage profiles suggested that PLE inhibits ICP1’s transition to rolling circle replication (Figure 2C). Since qPCR experiments indicated that the level of ICP1 replication was restored when PLE replication was abolished, we last wanted to determine if ICP1’s change in replication mode was also restored when PLE replication was abolished. Therefore we performed deep sequencing of total DNA during an ICP1 infection time course in PLE Δ*repA* and quantified and mapped coverage from 8 and 16 minute post-infection samples. As expected and consistent with the qPCR results, the relative abundance and GPM of ICP1 did not differ from what we saw for the PLE(-) infection conditions (Figure 8A, Supplementary Table S4). As seen before, the coverage profile of ICP1 at 8 minutes post-infection shows that ICP1 uses a bidirectional mode of replication at that time point. Interestingly, while loss of PLE replication restored ICP1 copy, abundance across the ICP1 genome matched neither the PLE(-) nor WT PLE(+) culture conditions at 16 minutes post-infection. The highest abundance of reads was shifted near to the end of ICP1’s annotated genome, as in the PLE(-) infection, but the gradual decrease in reads from this point was bidirectional rather than unidirectional (Figure 8B compare with Figure 2C). This reveals that PLE has some capacity to act on ICP1 replication even when the PLE is not replicating, and suggests that PLE may prevent linearization of ICP1’s genome.

**Figure 8.**
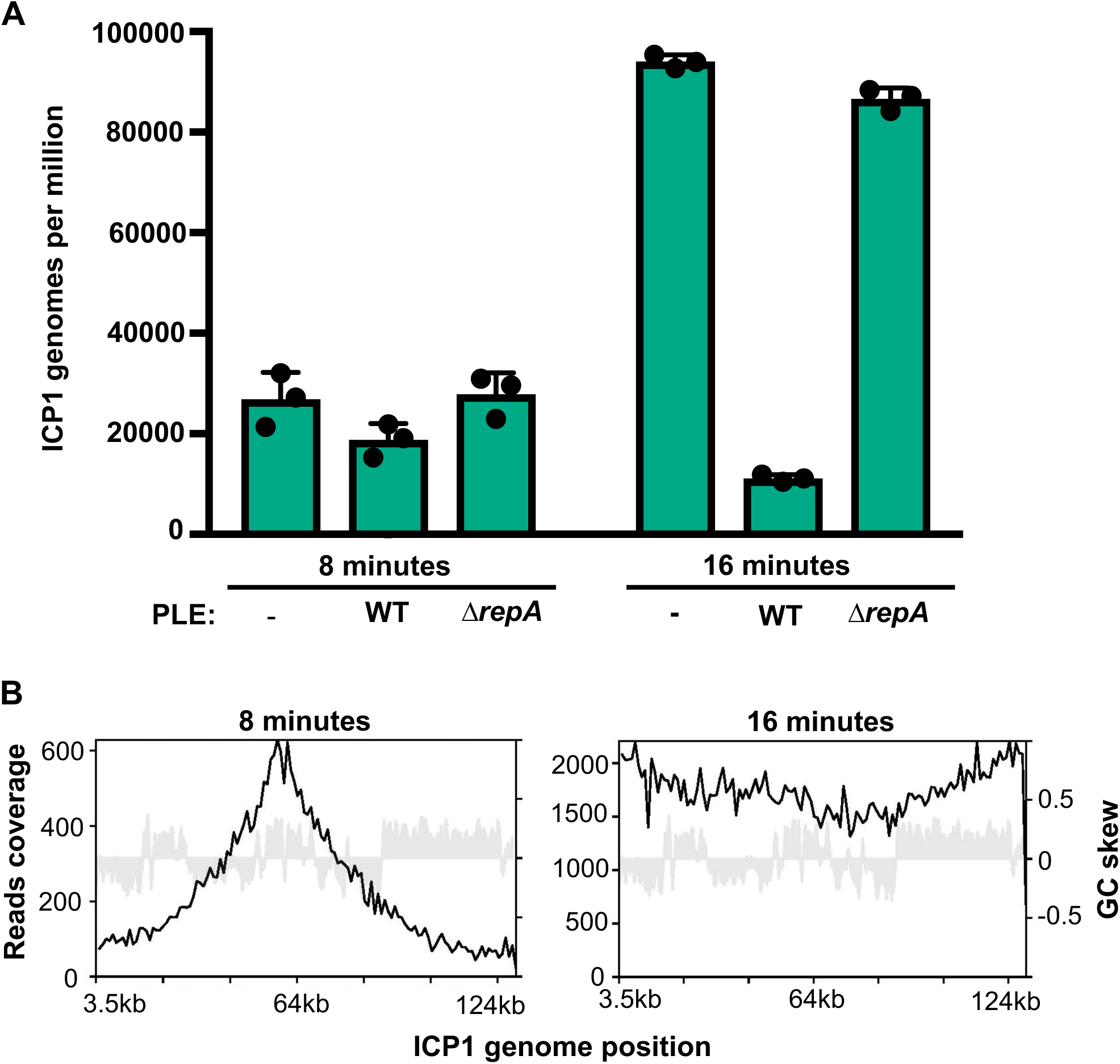
Non-replicating PLE still alters ICP1’s replication profile. (**A**) Genomes per million of ICP1 in PLE(-), PLE(+), and PLE Δ*repA* cultures at 8 and 16 minutes post-infection. Values shown are the means of three biological replicates. (**B**) Representative reads coverage profile of ICP1’s genome in Δ*repA* PLE infection at 8 minutes (left) and 16 minutes (right) post-infection. Coverage maps from replicate experiments are shown in Supplementary Figure S11.

## DISCUSSION

Here, we have identified the key constituents of PLE replication and evaluated their importance for ICP1 restriction. Since PLE’s genetic material outnumbers ICP1’s by 16 minutes post-infection, it is easy to imagine that reduced ICP1 copy, coupled with the presence of a highly abundant competing genome could severely hamper ICP1 packaging. Still, neither a decrease in ICP1 copies, nor a high abundance of PLE copies is necessary for PLE’s anti-phage activity, indicating that PLE has other mechanisms for restricting ICP1. Our results indicate that one of these mechanisms may still be centered around ICP1’s replication even if it does not decrease the overall ICP1 copy. The coverage profiles of ICP1 suggests that it undergoes a transition in replication mode from bidirectional replication to rolling circle replication, and that PLE impedes this transition even when PLE is unable to replicate (Supplementary Figure S12). A number of well characterized phages are known to transition from bidirectional theta to rolling circle replication over the course of infection (34). This transition linearizes and concatenates the phage genome, and concatemeric DNA serves as the packaging substrate for most tailed phage (43). If PLE Δ*repA* still prevents ICP1 from replicating via a rolling circle mechanism, this could severely impair ICP1’s ability to package its genome. PhageTerm analysis suggests that the ICP1 pac site is proximal to where we predict the rolling circle replication origin, potentially linking rolling circle replication and genome packaging. It is conceivable that the blunt terminus generated by the first round of rolling circle replication could act as a recognition site for the ICP1 terminase, which would then package the concatemeric genome in a headful fashion. Additionally, if a loss of genome linearization is not sufficient to prevent ICP1 particle production, it could act synergistically with other anti-packaging mechanisms such as the capsid hijacking observed in SaPIs and P4.

The precise relationship between ICP1’s and the midiPLE’s (and by extension PLE’s) DNA replication remains unclear. Specifically, it is unclear if replication of the midiPLE does not interfere with ICP1’s replication because the midiPLE is unable to replicate to the same level as PLE, or if the midiPLE is unable to reach as high of a copy level because ICP1 replication is unperturbed. Further work will be needed to identify factors that act on ICP1 replication without increasing PLE copy.

A major standing question for PLE replication is what replication machinery PLE is recruiting to the ori. Replication proteins for SaPIs and P4 possess primase and helicase activity (21–23) but PLE RepA is not predicted to possess either. How RepA_N proteins initiate replication remains unknown, and given that the C-terminal domain of these proteins are genera specific, it is likely that RepA_N proteins in different bacteria will recruit different machinery. Unlike the helper phages of SaPIs and P4, ICP1 possesses its own DNA polymerase, and it is not clear if *V. cholerae’s* replication machinery remains functional during ICP1 infection. Since PLE requires ICP1 infection to replicate even when RepA is ectopically expressed, we anticipate that PLE replication requires ICP1 gene products to replicate. ICP1 encodes a Pol-I type DNA polymerase and a helicase-primase like the *E. coli* phage T7 (16, 34, 44). The T7 replisome is one of the best characterized replisomes and is simpler in its components than most other replication complexes. Only four proteins are needed to reconstitute T7’s replisome *in vitro* (44): a DNA polymerase; host encoded thioredoxin, which acts as a processivity factor and prevents the aggregation of the cysteine-rich DNA polymerase; a helicase-primase, which in addition to possessing both helicase and primase activity has single stranded DNA binding activity and recruits the DNA polymerase; and a single stranded DNA binding protein that aids in replisome assembly and is necessary for lagging strand synthesis. To initiate PLE replication, RepA may recruit ICP1’s helicase-primase, or an accessory factor, such as a helicase or single strand binding protein, that remodels the ori in such a way that the helicase-primase will bind on its own and drive replisome assembly.

A third potential target for RepA would be an RNA polymerase. To initiate replication *in vivo*, T7 uses its self-encoded RNA polymerase (44). The primary and secondary T7 origins of replication are proximal to T7 RNA polymerase promoters. It is thought that the T7 RNA pol’s helicase activity provides free single stranded DNA for helicase-primase binding, and the T7 RNA polymerase lays the initial leading strand primer. A previous survey compared 15 divergent phages with T7-like replisomes, and 5 of the 15 were not predicted to encode their own RNA pol (34). Similarly, ICP1 does not encode an obvious RNA polymerase. Many phage encode factors for redirecting host RNA polymerase (34). For example, T4 is able to redirect the host RNA polymerase and uses it for initiation at replication origins (45), showing that transcription coupled replication can be employed by phage with divergent replisome types. If PLE uses ICP1’s DNA polymerase, PLE may need to recruit RNA polymerase to its ori for replication priming.

It is surprising that PLE encodes a RepA_N family initiator given their rarity among Gram-negative bacteria. Of the 742 RepA_N family proteins annotated in the pfam database, 723 belong either to Firmicutes species or bacteriophage that infect them (Pfam: Family:RepA_N (PF06970)). Only two RepA_N family sequences had been identified in proteobacteria, both of them in the group Burkholderiales. Interestingly, RepA is not the only PLE encoded gene that belongs to a family that is underrepresented in Gram-negative bacteria. The PLE integrase responsible for excision and integration into the host chromosome is a large serine recombinase (24) another protein type rarely found in Gram-negative bacteria and phage (46). Though PLEs lack any detectable homology to other known satellites, the presence of a large serine recombinase and RepA_N initiator in PLEs raises the possibility of recent inter-phyla gene transfer or deep evolutionary roots for PLEs.

Previously, it was noted that the chromosomally encoded RepA_N family proteins are linked to tyrosine and serine recombinases (40). The authors speculated that these genes were located on conjugative transposons and that the RepA_N had acquired new activities to facilitate transfer, since conjugative transposons, unlike plasmids or phages, do not need to replicate independently of the chromosome. An equally plausible explanation is that these recombinases and RepA_N genes are encoded by cryptic bacteriophage satellites. Supporting this possibility, a *Clostridium difficile* conjugative transposon encoding both a serine recombinase and a RepA_N initiator, as well as erythromycin resistance, was found to be transduced by a phage at a higher frequency of transfer than could be achieved by filter mating (47). This suggests that the boundary between viral satellite and conjugative element may not always be well defined, and individual elements may have some flexibility in their routes of mobilization. Since satellites typically do not encode their own structural genes, there is little to distinguish them from transposons or conjugative elements when one is making sequence based predictions. We anticipate that bacteriophage satellites will be found to be far more common than currently appreciated. Characterization of the PLE offers a window into these fascinating entities that shape the lives of their bacterial, and viral, hosts.

## DATA AVAILABILITY

During review a direct link to access the data has been provided. Prior to publication, the sequencing data from phage infected cells generated in this study will be deposited in the Sequence Read Archive database under accession codes TBD.

## SUPPLEMENTARY DATA STATEMENT

Supplementary Data are included.

## ACKNOWLEDGMENT

We thank Brendan O’Hara and Stephanie Hays for the construction of several strains used in this study, as well as Stephanie Hays for preparing the sequencing libraries and helpful advice for data presentation. We thank members of the Seed lab for useful discussion and feedback, and the Komeili lab for use of their AKTA for protein purification. We would also like to thank the University of California, Berkeley QB3 Core Facility for assistance with whole genome sequencing.

---

**Supplementary Figure 1.** ICP1 PLE(-) maps. Reads coverage profiles across the ICP1 genome for individual samples from replicate PLE(-) infection time course experiments.

**Supplementary Figure 2.** ICP1 PLE(+) maps. Reads coverage profiles across the ICP1 genome for individual samples from replicate PLE(+) infection time course experiments.

**Supplementary Figure 3.** Reads coverage profiles across the PLE genome for individual samples from replicate ICP1/PLE(+) infection time course experiments.

**Supplementary Figure 4.** ICP1 is predicted to use a headful packaging mechanism dependent on a pac site as determined by PhageTerm analysis. (**A**) PhageTerm schematic showing the predicted packaging mode of ICP1. ICP1 DNA is packaged into capsids using a headful mechanism from a distinct original site on the phage genome. (**B**) A zoomed in view of ICP1’s terminus with whole genome coverage plotted. (**C**) Plots of reads start position coverage divided by whole coverage along the entire ICP1 genome. The (+) strand (left) and (-) strand (right) are plotted separately.

**Supplementary Figure 5.** Repeat sequences found within the PLE noncoding region. Mismatches are shown in light gray, asterisks represent conserved sequence. Repeats 1 and 2 are interspersed with each other across a 528bp region (Figure 3B). The repeat 3 sequences are proximal to each other separated by 5 and 14bp, and the repeat 4 sequences are contiguous.

**Supplementary Figure 6.** Overlay of ribbon diagrams for the crystal structures of the NTD of dimers of PLE RepA (light blue) and pSK41 (dark grey) (RMSD = 4.197942 over 184 residues).

**Supplementary Figure 7.** Electrostatic profiles for PLE RepA (**A**), pTZ2162 RepA (**B**) pSK41 (**C**), and *B. subtilis* DnaD (**D**), N-terminal domain dimers. Positive (blue) and negative (red) charges are indicated on the surface.

**Supplementary Figure 8.** Replicates of an electrophoretic mobility shift assay using probes from the PLE noncoding region. RepA binding was tested for probes containing the repeat 3 or repeat 4 sequence from the PLE *NCR3*, as well as a probe matching the repeat 1 and repeat 2 sequences in *NCR2*.

**Supplementary Figure 9.** PLE replication aids transduction. Transduction units per mL produced from ICP1 infection of Δ*repA* PLE complemented with *repA* or an empty vector control. Quantification limit (QL) = 200 TU/mL.

**Supplementary Figure 10.** Spot assay showing ICP1 susceptibility for *V. cholerae* harboring the midiPLE complemented with an empty vector or *repA*.

**Supplementary Figure 11.** Reads coverage plots across the ICP1 genome for individual samples from replicate Δ*repA* PLE infection time course experiments.

**Supplementary Figure 12.** Model of PLE interference of ICP1 replication. ICP1 begins replication through a bidirectional theta mechanism before switching to rolling circle replication (RCR). RCR linearizes the ICP1 genome so that it can be packaged into capsids. The PLE is able to block ICP1 linearization without replicating. PLE inhibition of ICP1 copy increase is dependent on PLE replication.

