## Supplementary figures and tables for "Genome replication dynamics of a bacteriophage and its satellite reveal strategies for parasitism and viral restriction"

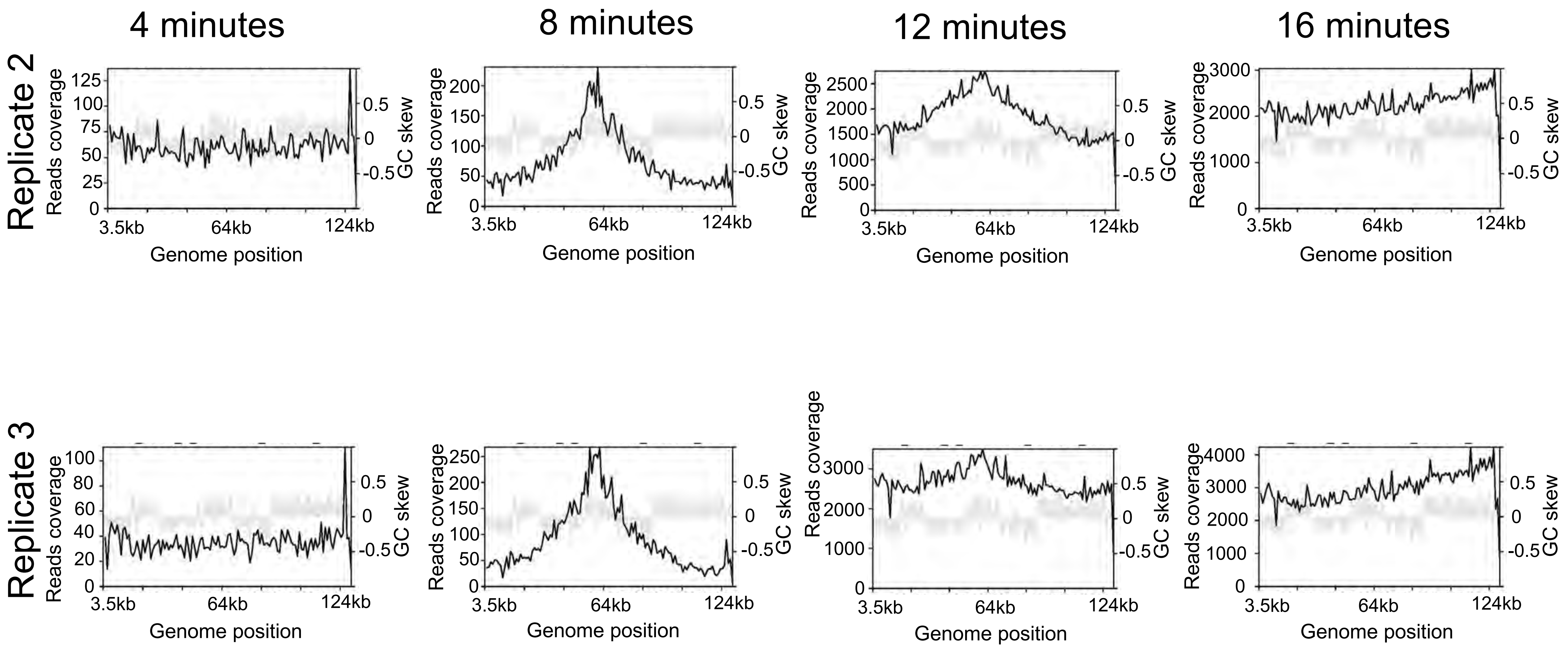


**Supplementary Figure 1.** ICP1 PLE(-) maps. Reads coverage profiles across the ICP1 genome for individual samples from replicate PLE(-) infection time course experiments.

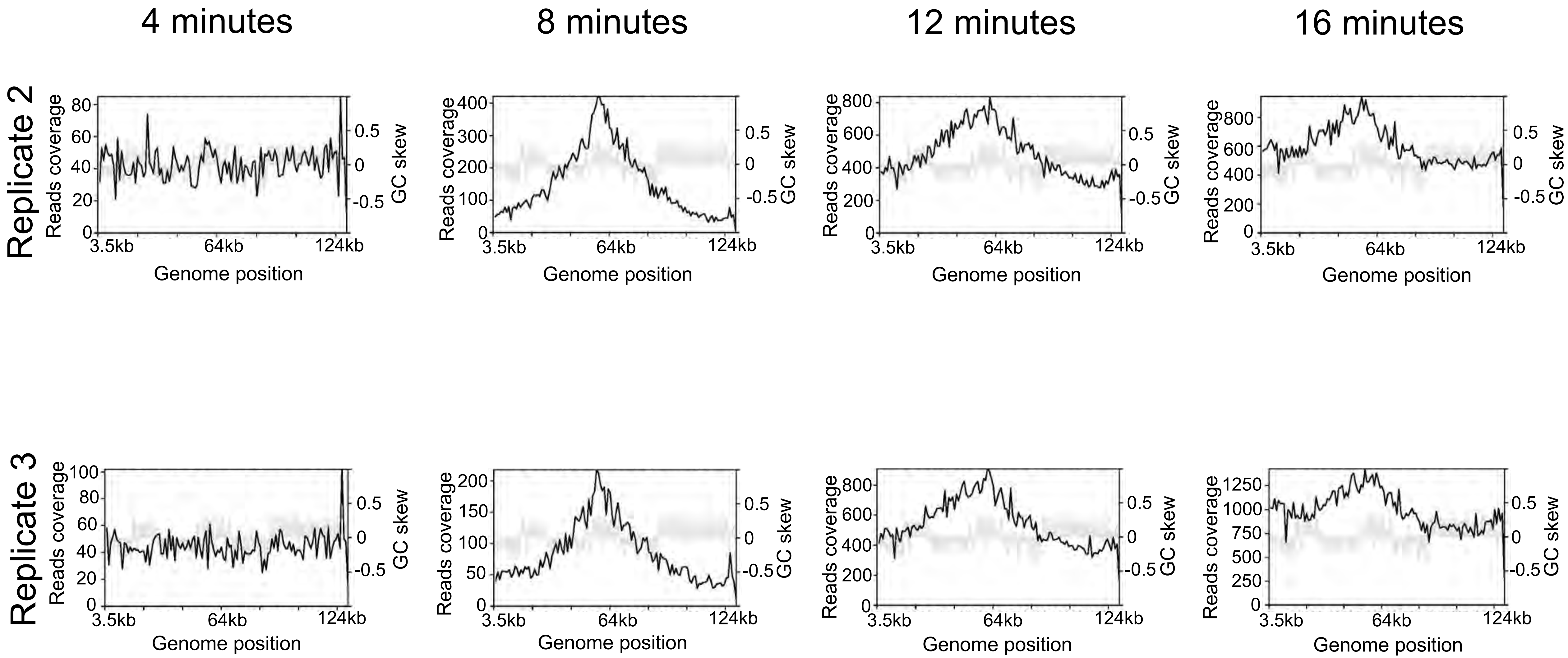


**Supplementary Figure 2.** ICP1 PLE(+) maps. Reads coverage profiles across the ICP1 genome for individual samples from replicate PLE(+) infection time course experiments.

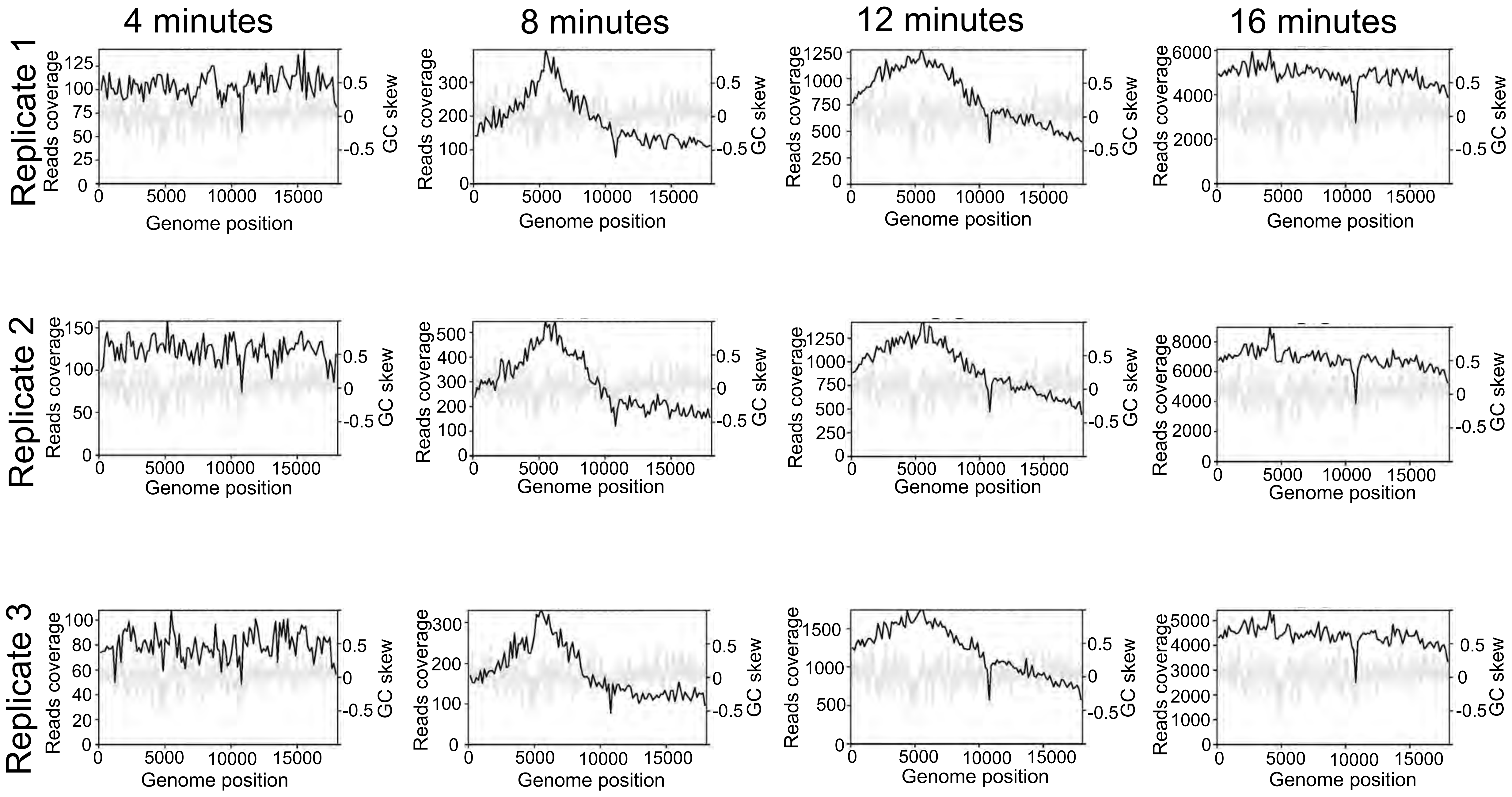


**Supplementary Figure 3.** Reads coverage profiles across the PLE genome for individual samples from replicate ICP1/PLE(+) infection time course experiments.

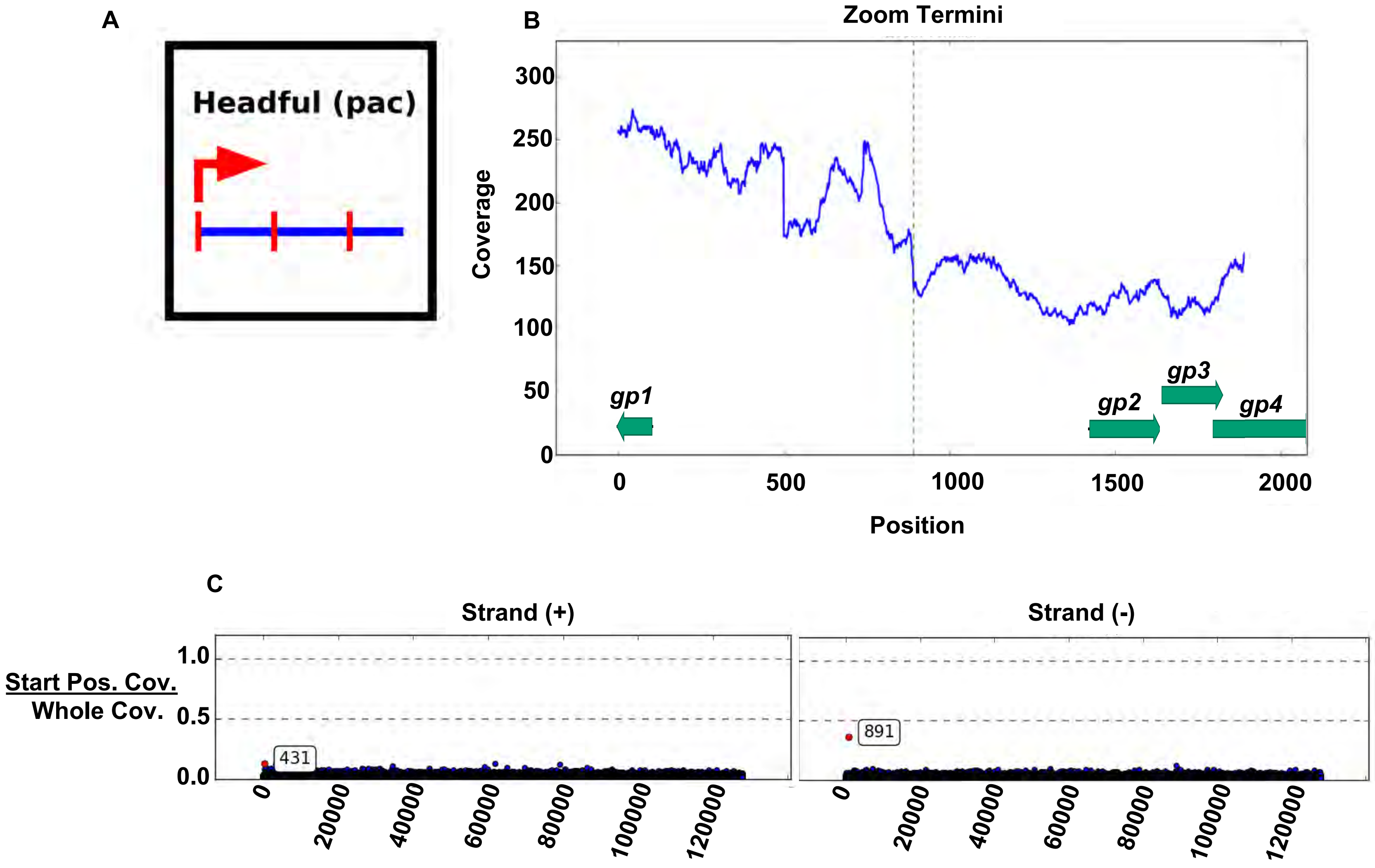


**Supplementary Figure 4.** ICP1 is predicted to use a headful packaging mechanism dependent on a pac site as determined by PhageTerm analysis. (**A**) PhageTerm schematic showing the predicted packaging mode of ICP1. ICP1 DNA is packaged into capsids using a headful mechanism from a distinct original site on the phage genome. (**B**) A zoomed in view of ICP1's terminus with whole genome coverage plotted. (**C**) Plots of reads start position coverage divided by whole coverage along the entire ICP1 genome. The (+) strand (left) and (-) strand (right) are plotted separately.

**Repeat 1**

```
CAGAACGTCATTTAACGCATCTTAT-CACCACCTTAATA
CAGAACGTCATTTAACGCATTTTACGCACCACCCTAATA
******************** ***  ******* *****
```

**Repeat 2**

```
ACTTATACGTTAGTATTACTGACGTTAGTATTACCCCCA
ACTTA--CGTTAGTATAACTTACGTTAGTA-TACCCTCA
*****  ********* *** ********* ***** **
```

**Repeat 3**

```
TTTATAGTTAGTGGGATGATTTTCATACCTATAAA
TTTATAT------GGATGATTTTCAACCCTATAAA
TTTATAG------GTATGATTTTCAGCCCTATAAA
******       * **********  ********
```

**Repeat 4**

```
AAGTGAGACACCTTATGGTAGTTC
AAGGTAGACACCTTATGGTAGTTC
AAGGTAGACACCTTTTGGTAG---
***  ********* ******
```

**Supplementary Figure 5.** Repeat sequences found within the PLE noncoding region. Mismatches are shown in light gray, asterisks represent conserved sequence. Repeats 1 and 2 are interspersed with each other across a 528bp region (Figure 3B). The repeat 3 sequences are proximal to each other separated by 5 and 14bp, and the repeat 4 sequences are contiguous.

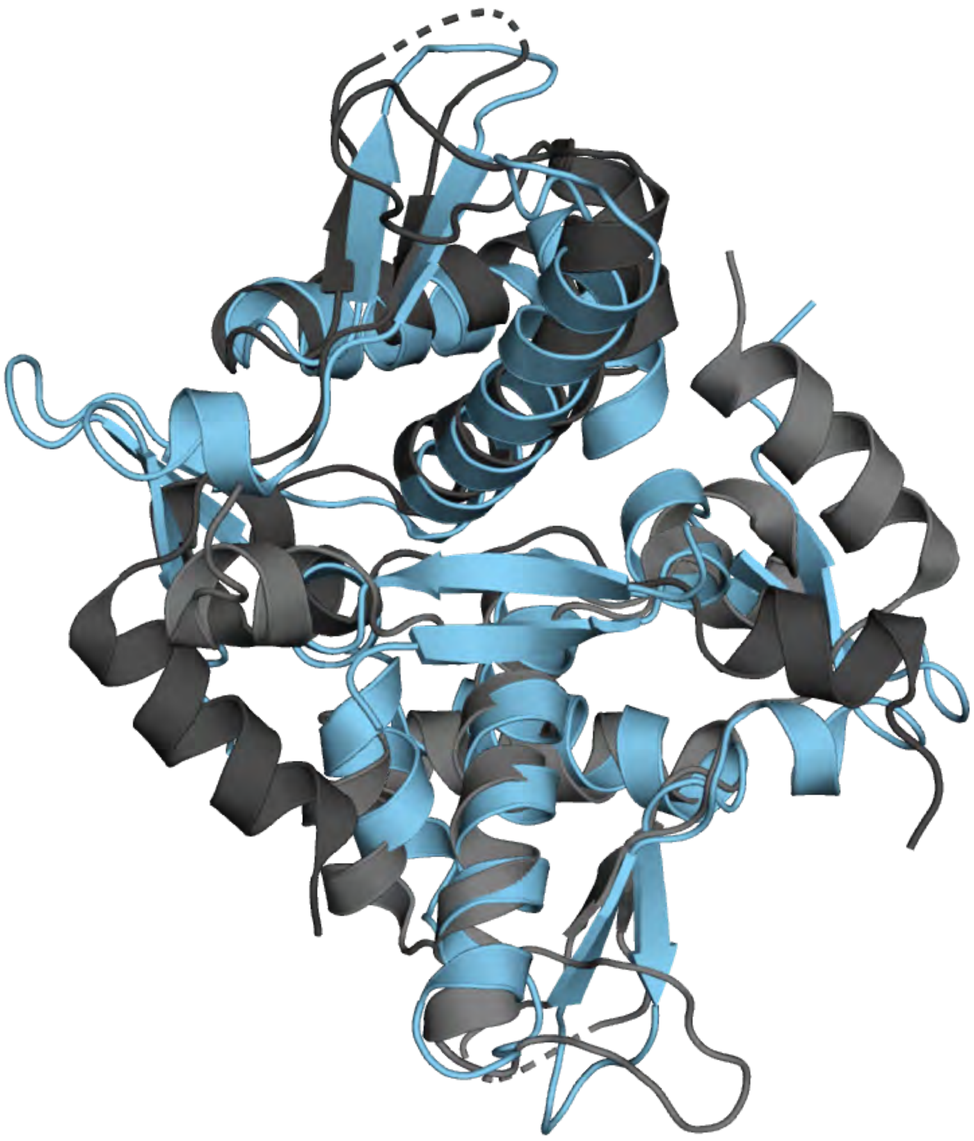

**Supplementary Figure 6.** Overlay of ribbon diagrams for the crystal structures of the NTD of dimers of PLE RepA (light blue) and pSK41 (dark grey) (RMSD = 4.197942 over 184 residues).

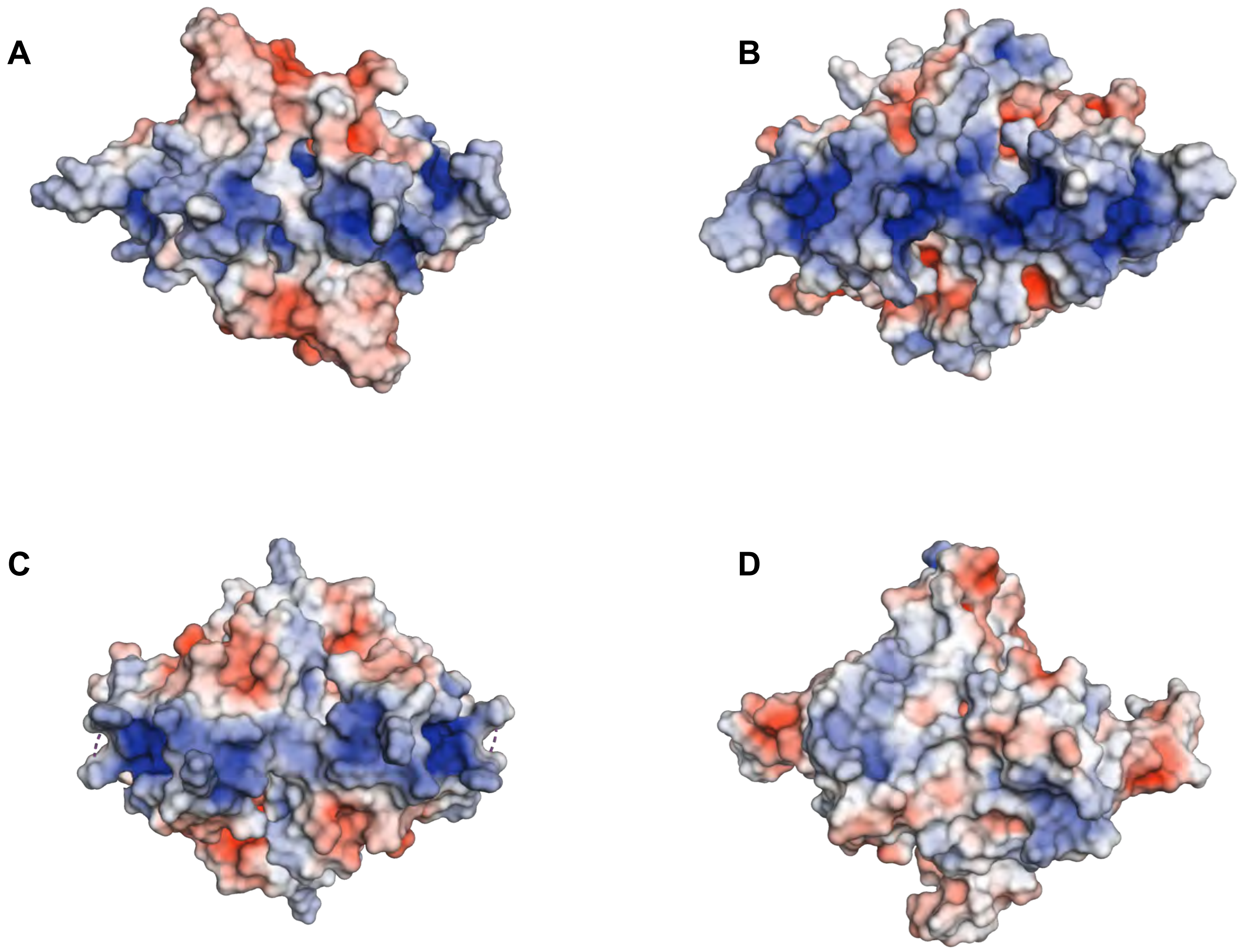


**Supplementary Figure 7.** Electrostatic profiles for PLE RepA (**A**), pTZ2162 RepA (**B**) pSK41 (**C**), and *B. subtilis* DnaD (**D**), N-terminal domain dimers. Positive (blue) and negative (red) charges are indicated on the surface.

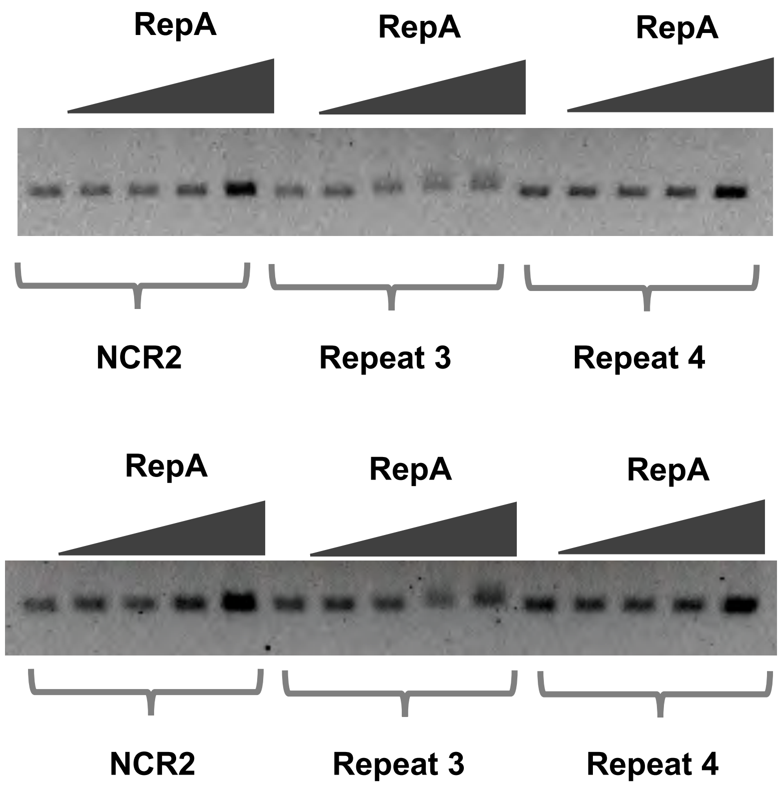


**Supplementary Figure 8.** Replicates of an electrophoretic mobility shift assay using probes from the PLE noncoding region. RepA binding was tested for probes containing the repeat 3 or repeat 4 sequence from the PLE *NCR3*, as well as a probe matching the repeat 1 and repeat 2 sequences deleted in the Δ*NRC2* mutant.

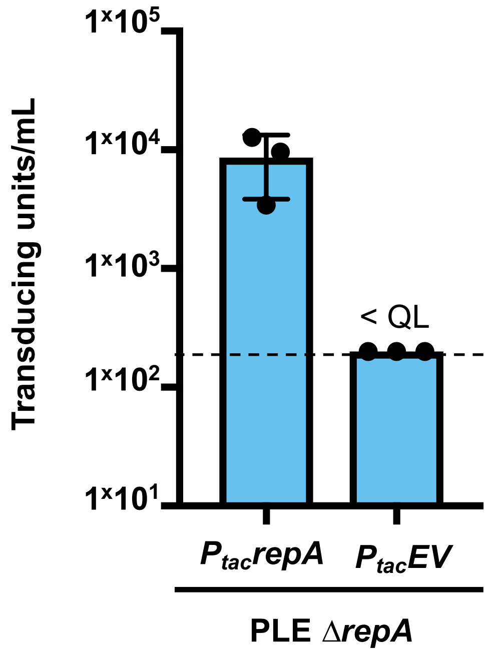


**Supplementary Figure 9.** PLE replication aids transduction. Transduction units per mL produced from ICP1 infection of Δ*repA* PLE complemented with *repA* or an empty vector control. Quantification limit (QL ) = 200 TU/mL.

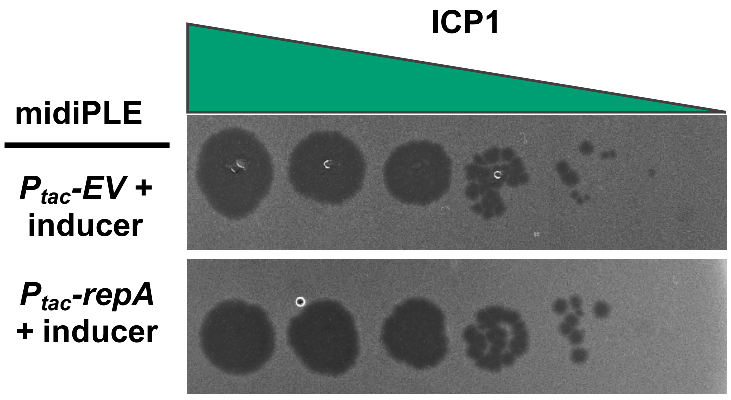


**Supplementary Figure 10.** Spot assay showing ICP1 susceptibility for *V. cholerae* harboring the midiPLE complemented with an empty vector or *repA*.

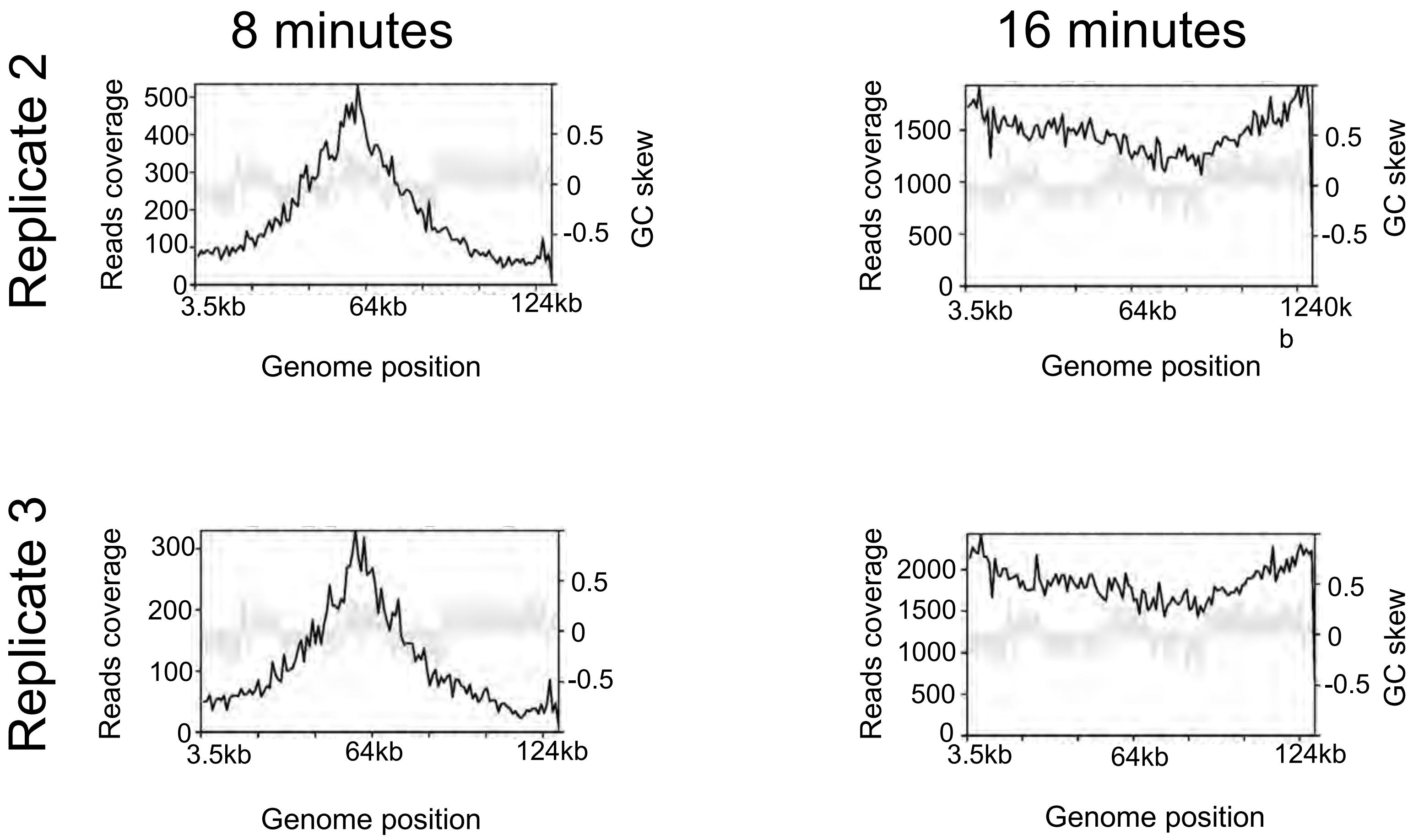


**Supplementary Figure 11.** Reads coverage plots across the ICP1 genome for individual samples from replicate Δ*repA* PLE infection time course experiments.

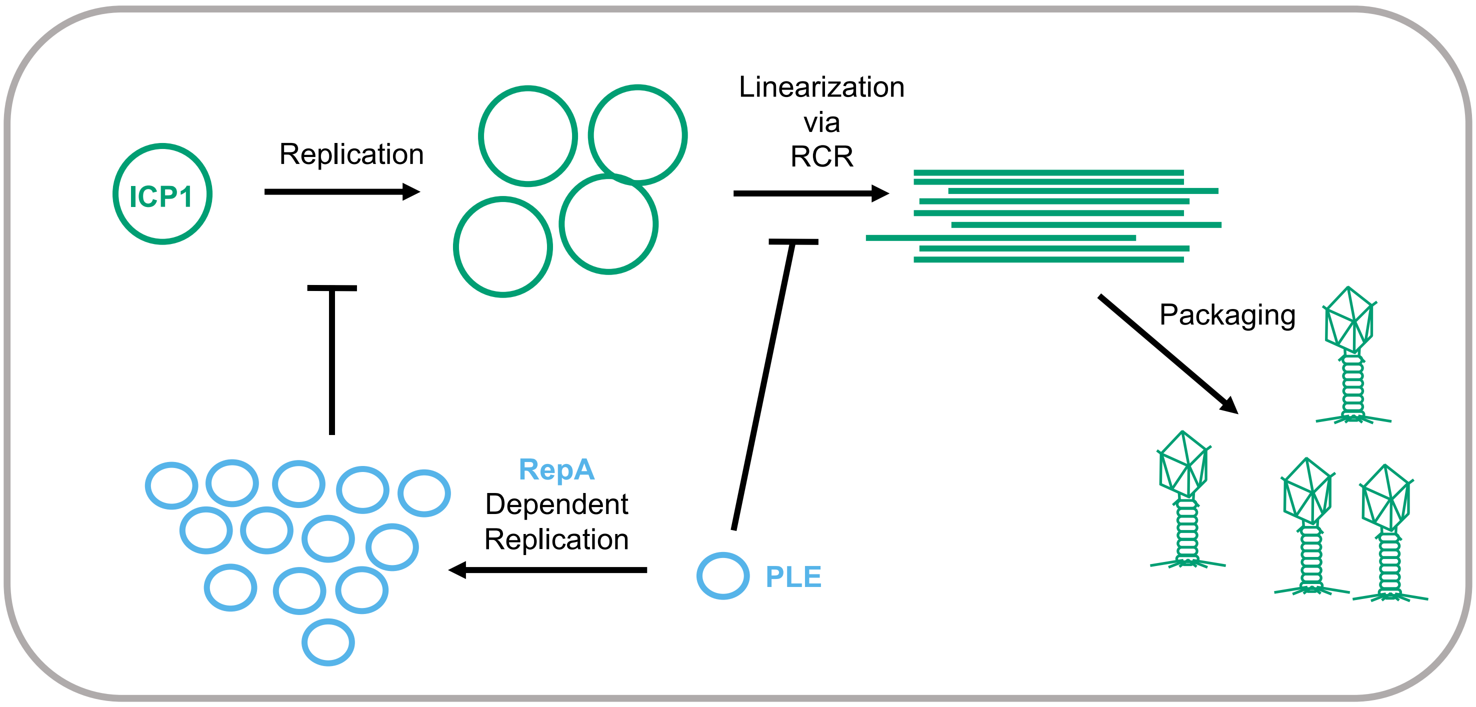


**Supplementary Figure 12.** Model of PLE interference of ICP1 replication. ICP1 begins replication through a bidirectional theta mechanism before switching to rolling circle replication (RCR). RCR linearizes the ICP1 genome so that it can be packaged into capsids. The PLE is able to block ICP1 linearization without replicating. PLE inhibition of ICP1 copy increase is dependent on PLE replication.

**Supplementary Table S1: Strains used in this study**

| Strain | Description* | Source |
|---|---|---|
| KDS6 | *V. cholerae* O1, El Tor biotype | Lab collection |
| KDS36 | *V. cholerae* E7946 containing PLE1 | (1) |
| KDS153 | *V. cholerae* E7946 PLE1 ΔORF7::frt, KanR | This study |
| KDS154 | *V. cholerae* E7946 PLE1 ΔORF8::frt, KanR | This study |
| KDS155 | *V. cholerae* E7946 PLE1 ΔORF9::frt, KanR | This study |
| KDS156 | *V. cholerae* E7946 PLE1 ΔORF10::frt, KanR | This study |
| KDS157 | *V. cholerae* E7946 PLE1 *ΔrepA*::frt, KanR | This study |
| KDS158 | *V. cholerae* E7946 PLE1 ΔORF12::frt, KanR | This study |
| KDS159 | *V. cholerae* E7946 PLE1 ΔORF12.1::frt, KanR | This study |
| KDS160 | *V. cholerae* E7946 PLE1 ΔORF13::frt, KanR | This study |
| KDS161 | *V. cholerae* E7946 PLE1 ΔORF14::frt, KanR | This study |
| KDS181 | *V. cholerae* E7946 PLE1 *Δint*::Spec-frt | (2) |
| KDS182 | *V. cholerae* E7946 PLE1 ΔORFs2-5::Spec-frt | (2) |
| KDS183 | *V. cholerae* E7946 PLE1 ΔORFs7-14::Spec-frt | (2) |
| KDS184 | *V. cholerae* E7946 PLE1 ΔORFs15-20::Spec-frt | (2) |
| KDS185 | *V. cholerae* E7946 PLE1 ΔORFs21-23::Spec-frt | (2) |
| KDS228 | *V. cholerae* E7946 *ΔlacZ*::SpecR | (1) |
| KDS229 | *V. cholerae* E7946 PLE1 *ΔrepA*::frt, *ΔlacZ*::$P_{tac}$-*repA*, KanR, SpecR (RepA chromosomal expression construct in PLE *ΔrepA*) | This study |
| KDS230 | *V. cholerae* E7946 PLE1 *ΔrepA*::frt, *ΔlacZ*:: $P_{tac}$ *EV*, KanR, SpecR (Empty chromosomal expression construct in PLE *ΔrepA* ) | This study |
| KDS231 | *V. cholerae* E7946 PLE1 *ΔrepA*::frt, $P_{tac}$-*repA*, KanR, CmR (Plasmid RepA expression construct) | This study |
| KDS232 | *V. cholerae* E7946 midiPLE, $P_{tac}$-*repA*, KanR, SpecR (RepA plasmid expression construct in strain with midiPLE) | This study |
| KDS233 | *V. cholerae* E7946 PLE1 *ΔrepA*::frt, $P_{tac}$ *EV*, KanR, CmR (Empty plasmid expression construct in PLE *ΔrepA* ) | This study |
| KDS234 | *V. cholerae* E7946 midiPLE, $P_{tac}$ *EV*, KanR, CmR (Empty plasmid expression construct in strain with midiPLE) | This study |
| KDS235 | *V. cholerae* E7946 PLE1 ΔNCR1, KanR | This study |
| KDS236 | *V. cholerae* E7946 PLE1 ΔNCR2::frt, KanR | This study |
| KDS237 | *V. cholerae* E7946 PLE1 ΔNCR3::frt, KanR | This study |
| KDS238 | *E. coli* BL21 pE-SUMO-RepA. Vector to express 6xHisSumo-fusion protein, fused to N-terminus of RepA | This study |
| ICP1 | ICP1_2006_E ΔCRISPR ΔCas2-3 | (2) |

* KanR = Kanamycin resistance cassette, SpecR = Spectinomycin resistance cassette.

**Supplementary Table S2: Primers used in this study**

| Primer | Sequence | Purpose | Source |
|---|---|---|---|
| zac14 | AGGGTTTGAGTGCGATTACG | qPCR PLE | (1) |
| zac15 | TGAGGTTTTACCACCTTTTGC | qPCR PLE | (1) |
| zac68 | CTGAATCGCCCTACCCGTAC | qPCR ICP1 | (1) |
| zac69 | GTGAACCAACCTTTGTCGCC | qPCR ICP1 | (1) |
| zac109 | CGCCAAACCAACAAGACAGG | qPCR PLE ΔNCR | This study |
| zac110 | CCCCAAGATCAACCACCTCC | qPCR PLE ΔNCR | This study |
| zac192 | CGAAGCGTGTTCTTTATAGTTAGTGG | F primer Repeat 3 probe | This study |
| zac193 | CTGTAACACTGTATGAATTTATAGGGC | R primer Repeat 3 probe | This study |
| zac194 | CCTAGAATATCAATCGCTTACCAG | F primer Repeat 4 probe | This study |
| zac195 | GCAACATATGATTGTGTGATGCC | R primer Repeat 4 probe | This study |
| zac196 | CCACTTATACGTTAGTATTACTGACG | F primer Repeat 1+2 | This study |
| zac197 | GGTATACTAATGTATTAGGGTGGTGC | R primer Repeat 1+2 | This study |

**Supplementary Table S3: Proportional reads abundance for ICP1, PLE, and the *V. cholerae* chromosomes relative to total sample reads over time course of ICP1 infection**

| | Percent reads per element of total reads per condition | | | | | | |
|---|---|---|---|---|---|---|---|
| | PLE(-) infection | | | PLE(+) infection | | | |
| Min[a] | VC I[b] | VC II | ICP1 | VC I | VC II | ICP1 | PLE |
| 4 | 79 ± 0. 27[c] | 20 ± 0.098 | 1.1 ± 0.17 | 79 ± 0.26 | 20 ± 0.15 | 1.1 ± 0.26 | 0. 32 ± 8.6E-04 |
| 8 | 78 ± 6.0 | 20 ± 0.097 | 2.2 ± 0.097 | 77 ± 0.57 | 19 ± 0.14 | 2.5 ± 0.61 | 0.67 ± 0.12 |
| 12 | 61 ± 5.9 | 15 ± 1.6 | 24 ± 7.4 | 65 ± 0.71 | 17 ± 0.17 | 14 ± 0.71 | 4.5 ± 0.29 |
| 16 | 41 ± 4.8 | 9.9 ± 1.2 | 49 ± 6.0 | 51 ± 1.3 | 14 ± 0.35 | 17 ± 0.69 | 19 ± 1.4 |

[a] time in minutes post-infection
[b] VC I is the *V. cholerae* large chromosome, VCI II is the *V. cholerae* small chromosome
[c] Values are the mean and standard deviation of three biological replicates

**Supplementary Table S4: Proportional reads abundance for ICP1, PLE, and the *V. cholerae* chromosomes relative to total sample reads over time course of ICP1 infection in PLE *ΔrepA***

| | Percent reads per element of total reads per condition | | | |
|---|---|---|---|---|
| Min[a] | VC I | VC II | ICP1 | PLE |
| 8 | 77 ± 0.56 | 19 ± 0.12 | 30 ± 0.62 | 0.31 ±0.013 |
| 16 | 44 ± 3.1 | 11 ± 0.8 | 45 ± 0.39 | 0.24 ± 8.5E -03 |

[a] time in minutes post-infection
[b] VC I is the *V. cholerae* large chromosome, VCI II is the *V. cholerae* small chromosome
[c] Values are the mean and standard deviation of three biological replicates
